# Src activation in lipid rafts confers epithelial cells with invasive potential to escape from apical extrusion during cell competition

**DOI:** 10.1101/2021.05.29.446275

**Authors:** Kentaro Kajiwara, Ping-Kuan Chen, Shunsuke Kon, Yasuyuki Fujita, Masato Okada

**Author notes:** Corresponding author Mailing address: Department of Oncogene Research, Research Institute for Microbial Diseases, Osaka University, Yamada-Oka 3-1, Suita 565-0871, Japan.

## Abstract

Abnormal/cancerous cells within healthy epithelial tissues undergo apical extrusion to protect against carcinogenesis, while they acquire invasive capacity once carcinogenesis progresses. However, the molecular mechanisms by which cancer cells escape from apical extrusion and invade surrounding tissues remain elusive. We found that during competition within epithelial cell layers, Src-transformed cells underwent basal delamination by Src activation within lipid rafts, whereas they were apically extruded when Src was outside of lipid rafts. Comparative analysis of contrasting phenotypes revealed that activation of the Src-STAT3-MMP axis through lipid rafts was required for basal delamination. CUB domain-containing protein 1 (CDCP1) was identified as an Src activating scaffold in lipid rafts, and its overexpression induced basal delamination. In renal cancer spheroids, CDCP1 promoted HGF-dependent invasion by activating the Src-STAT3-MMP axis. Overall, these results suggest that Src activation in lipid raft confers resistance to apical extrusion and invasive potential on epithelial cells to promote carcinogenesis.

## Introduction

The normal epithelial cell layer is maintained through various phenomena including turnover of harmful and suboptimal cells for homeostasis in healthy tissues. Oncogene-transformed mammalian cells are forcibly eliminated from the epithelial cell layer without involving the immune system^1-3^. Emerging transformed cells in normal cell layer are extruded by the surrounding normal cells in a non-cell-autonomous fashion, resulting in the maintenance of a healthy cell layer^4-7^. This phenomenon, known as apical extrusion, is a mode of “cell competition”, and is considered a maintenance system for healthy tissues in the initial phase of carcinogenesis^5, 8^. During cancer progression, such defence systems may be abrogated or overwhelmed by additional changes in cancer cells. However, little is known regarding how cancer cells escape this safeguard, expand their own area, and invade the surrounding normal tissues.

An oncogene product Src, a membrane-anchored tyrosine kinase, plays a pivotal role in regulating various cellular functions including cell adhesion, migration, and invasion^9^. Src is overexpressed and/or activated in various malignant cancers implying a crucial role in cancer progression. However, Src-transformed cells are eliminated from normal epithelial tissues by cell competition in invertebrate models using zebrafish embryos and *Drosophila* tissues, as well as in mammalian models^10-19^. Thus, Src showed dual functions depending on the cellular conditions. Previous studies using Src-activated MDCK type II cells have identified several cell competition-related proteins^3^. Filamin and vimentin accumulate in normal cells at the interface with Src-transformed cells^20^, whereas EPLIN accumulates in transformed cells^21^. Similar cellular events are also observed in oncogenic Ras-transformed cells^20, 21^. Cytoskeletal remodelling may also contribute to the apical extrusion of transformed cells from normal tissues^6, 12^. However, the mechanisms by which cell competition-related functions of Src are switched to malignancy-inducing functions in cancer cells are largely unknown.

Upon activation, Src is translocated between the cytosolic space and intracellular membrane to activate region-specific substrates^9^. Part of activated Src is translocated into sphingolipid/cholesterol-enriched lipid rafts through myristoylation to phosphorylate specific downstream signalling molecules^22-24^, indicating that Src function can be spatially regulated through association with lipid rafts. Src recruitment to lipid rafts is mediated by specific transmembrane scaffolds including Pag1/Cbp and CUB domain-containing protein 1 (CDCP1)^22, 25-28^. CDCP1 has two palmitoylation sites required for raft localisation and an Src-binding motif in the cytoplasmic domain, enabling Src activation in lipid rafts. Furthermore, CDCP1 associates with the HGF receptor Met, to facilitate Src-mediated STAT3 activation, which promotes the expression of matrix metalloproteases (MMPs) involved in extracellular matrix (ECM) remodelling^26^. Thus, spatial regulation of activated Src is important for sorting specific signal transductions, leading to the onset of physiological and pathological functions.

In this study, to address the molecular mechanisms by which Src-transformed cells escape from apical extrusion to acquire malignant properties, we analysed the behaviour of two types Src-transformed MDCK cells. Comparative analysis of these cells revealed that activation of the Src-STAT3-MMP axis through lipid rafts is required for basal delamination. We also show that CDCP1 induces basal invasion by Src activation in MDCK cells, and upregulation of CDCP1 promotes HGF-dependent invasion of renal cancer cells. These findings suggest that spatial activation of Src signalling in lipid raft confers malignant potential on epithelial cells to promote carcinogenesis by escaping from apical extrusion. This is the first report demonstrating a molecular mechanism for cell fate switching during cell competition.

## Results

### Src-transformed MDCK type I cells undergo basal delamination

To investigate the effects of Src transformation on epithelial cell competition, we introduced an inducible Src (EGFP-tagged) expression system into MDCK type I cells (Src-EGFP cells). After doxycycline (Dox) treatment, active Src was overexpressed (p-Src blot, pTyr418) and phosphorylated various intracellular proteins (pY1000 blot) (**Fig. 1a**). A kinase-defective mutant (Src-KD-EGFP) was used as a negative control. To monitor the behaviour of Src-transformed cells under cell competition condition, Src-EGFP cells were co-cultured with normal cells on a two-dimensional collagen gel. Before Dox treatment, Src-EGFP cells remained in the cell layer; overexpression of Src-EGFP, but not Src-KD-EGFP, induced cell delamination into the basal side of the cell layer (**Fig. 1b**). These phenomena were quantified based on the criteria defined by the vertical location of cells in the cell layer (**Fig. 1c**). In contrast, when only Src-EGFP cells were cultured, delamination was not observed even after Dox treatment (**Fig. 1b**). Furthermore, the efficacy of delamination in Src-EGFP cells was dependent on the density of normal cells and Src activation levels (**Supplementary Fig. 1a-d**). These results indicate that Src-induced basal delamination requires surrounding normal cells, which is a typical feature of epithelial cell competition. We also found that Src-EGFP cells adopted a round shape and quickly moved toward the apical side in the early phase of Src induction (∼12h), whereas they began to undergo delamination from 18 h after induction (**Fig. 1d, e**). These findings indicate that Src activation in MDCK type I cells induces basal delamination in the context of cell competition through multiple processes.

**Figure 1.**
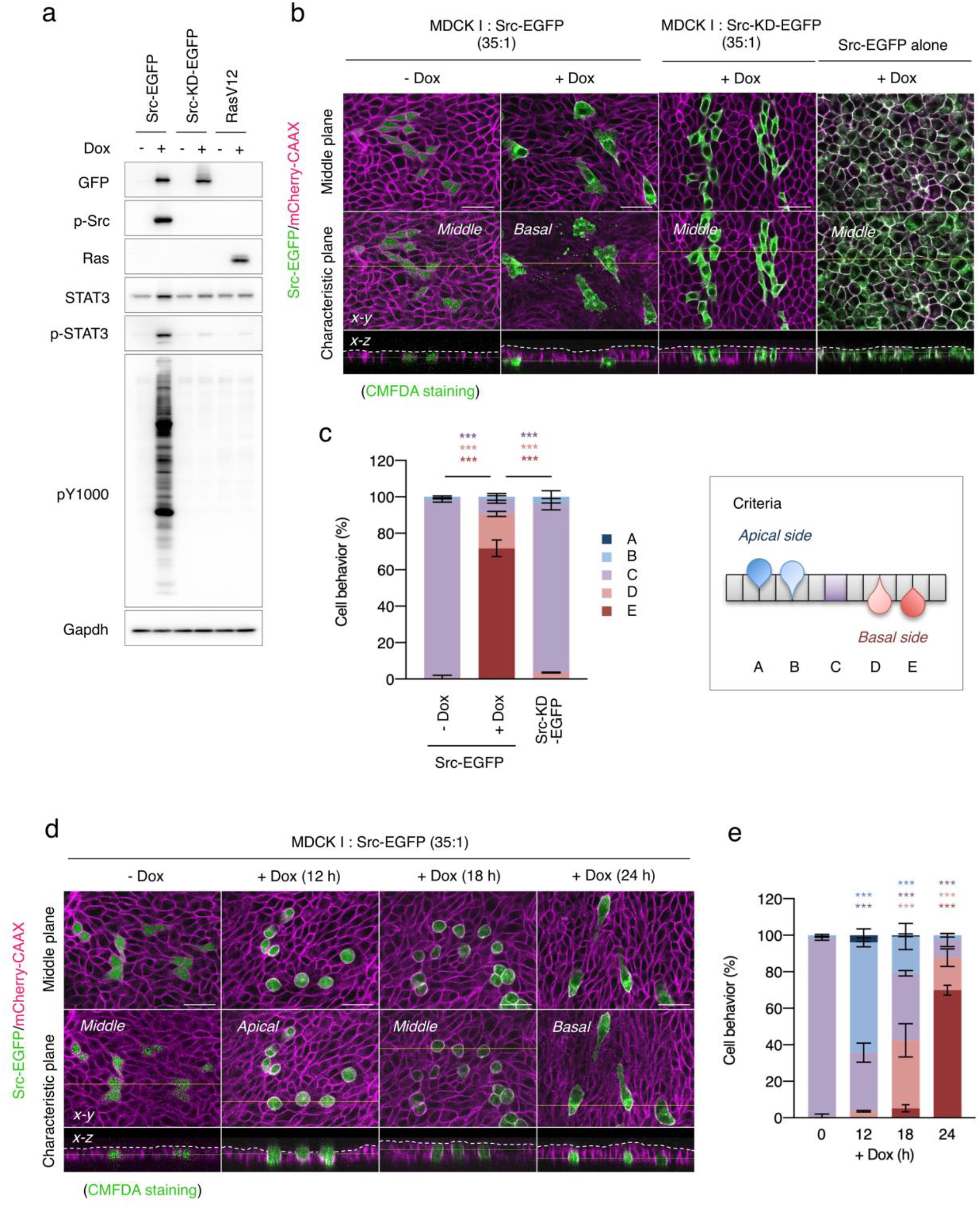
Src-transformed cells delaminate into the basal side of epithelial cell layer. (**a**) MDCK type I cells harbouring inducible expression system of Src-EGFP, Src-K297M-EGFP or RasV12 were grown in the presence or absence of 1 μg/ml doxycycline (Dox) for 24 h. Cell lysates were subjected to immunoblotting analysis using the indicated antibodies. (**b, c**) Type I cells harbouring Src-EGFP or Src-KD-EGFP were mixed with wild-type cells at a ratio of 35:1 and grown on collagen matrix in the presence or absence of 1 μg/ml Dox for 24 h. mCherry-CAAX is used as a cell membrane marker, and CMFDA is used as a fluorescein indicator of Src-EGFP harbouring cells in the absence of Dox. Cell behaviour was monitored (n > 100) and assessed by using criteria (c, right panel); A, apical extruded cell; B, apical extruding cell; C, staying cell; D, basal delaminating cell; E, basal delaminated cell. (**d, e**) Type I cells harbouring Src-EGFP were mixed with wild-type cells at a ratio of 35:1 and grown in the presence or absence of 1 μg/ml Dox for the indicated time periods. The mean ratios ± SDs were obtained from three independent experiments. ***, *P* < 0.001; two-way ANOVA was calculated compared with the Src-EGFP expressed cells (c) or non-treated cells (e). The scale bars indicate 50 μm.

However, our findings in MDCK type I cells were inconsistent with previous results obtained in other subtypes of MDCK type II cells^12, 20^. When Src-EGFP cells were co-cultured with normal cells in a three-dimensional collagen matrix to form mosaic cysts, Src-expressing MDCK type I cells underwent basal delamination (**Supplementary Fig. 1e**), as observed in two-dimensional cultures. However, Src-expressing MDCK type II cells were apically extruded and accumulated in the cyst lumen, as described previously^3, 5^. To examine whether induction of opposing phenotypes between MDCK type I and type II cells is unique to Src-transformed cells, we introduced oncogenic RasV12 in MDCK type I cells and performed a two-dimensional cell competition assay (**Supplementary Fig. 1f, g**). Consistent with previous studies in MDCK type II cells, RasV12-transformed cells were apically excluded even in MDCK type I cells. These results suggest that basal delamination is a unique feature of Src-transformed MDCK type I cells, and that comparative analysis between the two types of cells may provide mechanistic insights into the cell competition specifically induced by Src transformation.

### Focal adhesions are matured in Src-transformed MDCK type I cells

To address the mechanisms by which Src activation induces basal delamination in MDCK type I cells, we first examined the accumulation of an apical extrusion marker, EPLIN, identified in MDCK type II cells^21^. Immunofluorescence analysis revealed that EPLIN, a cytoskeleton-associated protein, was also accumulated in Src-transformed MDCK type I cells (**Supplementary Fig. 2a**), indicating that Src-induced basal delamination of MDCK type I cells shares some mechanisms observed in type II cells. Next, we observed the cellular events in Src-transformed cells undergoing basal delamination. Immunofluorescence indicated that activated Src was concentrated in focal adhesions facing the basement membrane, where phosphorylated FAK, filamentous actin, cortactin, and paxillin were also concentrated (**Supplementary Fig. 2b**). FAK inhibition by a specific inhibitor or by overexpression of dominant-negative FAK (FAK-DN) significantly attenuated Src-induced basal delamination (**Supplementary Fig. 2c, d**), indicating that Src-transformed cells delaminate through focal adhesion maturation. Furthermore, analysis using DQ-conjugated collagen, which can monitor the degradation of collagen matrix, revealed that Src-transformed delaminating cells have strong matrix degradation activity (**Supplementary Fig. 2e**). These results indicate that Src transformation promotes maturation of focal adhesions and delamination into the basement membrane through degradation of the extracellular matrix, suggesting that Src-transformed MDCK type I cells acquire invasive potential to escape apical extrusion.

### Lipid raft localisation is crucial for Src-induced basal delamination

To further identify the components required for Src-induced basal delamination, we performed inhibitor screening (**Supplementary Table 1**). Pretreatment with lipid metabolism inhibitors targeting the biosynthesis of sphingolipid, glycosphingolipid, fatty acids, and cholesterol, suppressed Src-induced basal delamination (**Fig. 2a**). These inhibitors suppress lipid synthesis and metabolism, leading to altered integrity of lipid rafts. As activated Src is translocated and concentrated into lipid rafts, Src distribution to lipid rafts may be involved in regulating Src-induced basal delamination. Indeed, activated Src was concentrated at the plasma membrane, where the lipid raft-component, flotillin1, was concentrated (**Fig. 2b**). Cholesterol levels in detergent-resistant membrane (DRM) fractions containing lipid raft components were gradually increased after Src overexpression (**Fig. 2c**). Furthermore, membrane cholesterol depletion by pretreatment with MβCD suppressed the basal delamination of Src-transformed cells, though some of cells were apically extruded (**Fig. 2a, d**). Comparative analysis of lipid raft Src localisation between MDCK type I and II cells showed that activated Src was immediately translocated into the DRM fractions in MDCK type I cells, whereas Src translocation proceeded significantly more slowly in MDCK type II cells than in type I cells (**Fig. 2e, f**). These results suggest that lipid raft Src localisation is involved in the induction of basal delamination.

**Figure 2.**
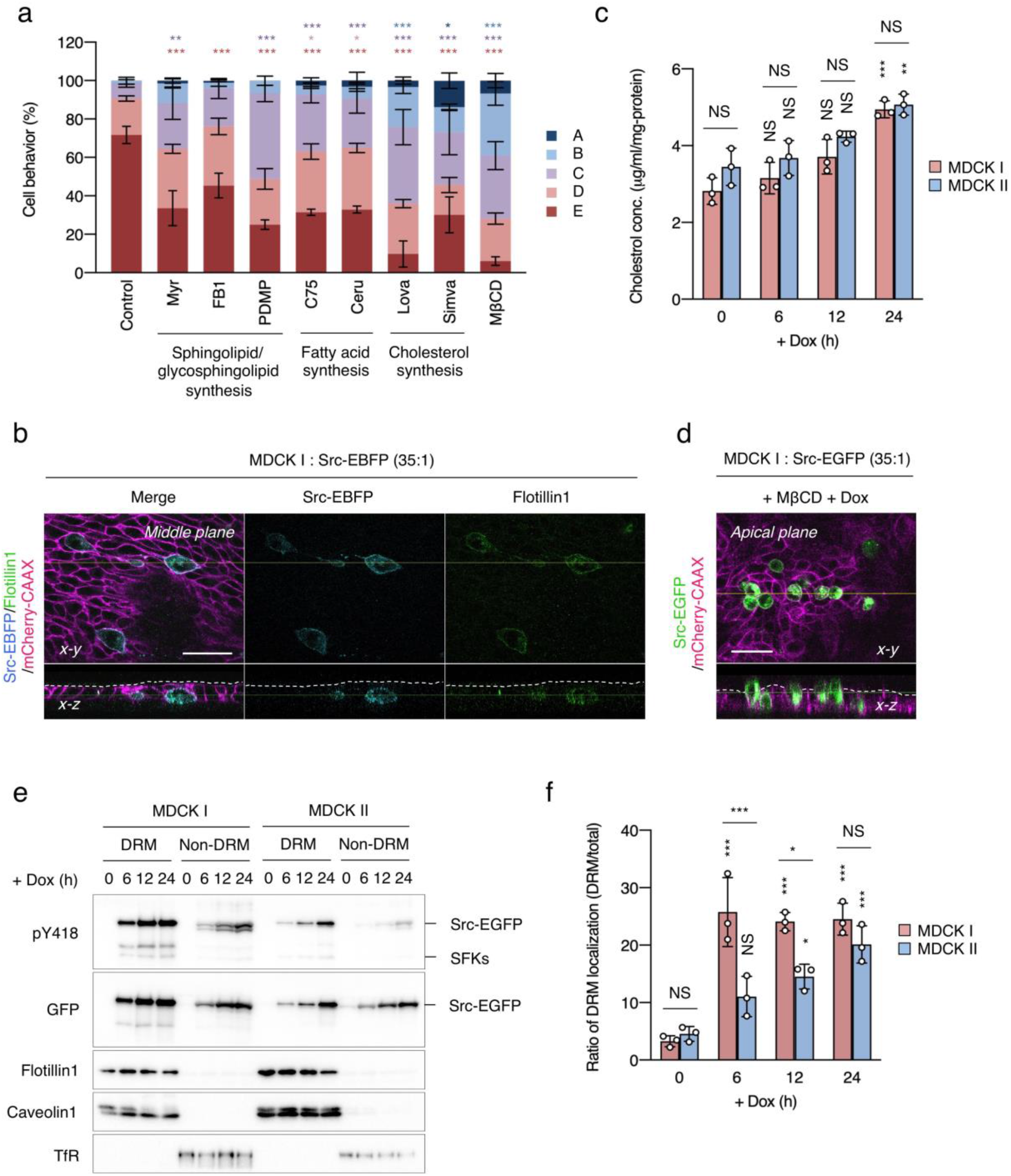
Lipid raft Src localisation is crucial for basal delamination. (**a**) MDCK type I cells harbouring Src-EGFP were mixed with wild-type cells at a ratio of 35:1 and preincubated with 200 nM myriocin (Myr), 200 nM fumonisin b1 (FB1), 400 nM D-threo-PDMP (PDMP), 400 nM C75, 400 nM cerulenin (Ceru), 400 nM lovastatin (Lova), 1.6 μM simvastatin (Simva), or 500 nM MβCD for 2 h and then incubated with Dox for 24 h. Cell behaviour was assessed by using the criteria (Fig. 1c). (**b**) Type I cells harbouring Src-EBFP were mixed with wild-type cells at a ratio of 35:1 and grown in the presence of Dox for 24 h. After fixation, flotillin1 was visualized with specific antibody and Alexa488-conjugated secondary antibody. (**c**) Type I and II harbouring Src-EGFP were grown in the presence of Dox for indicated time periods. Lysates from these cells were subjected to the DRM separation experiment to separate DRM (lipid raft) and non-DRM (non-lipid raft) fractions. The DRM fractions were subjected to cholesterol assay. (**d**) Type I cells harbouring Src-EGFP were mixed with wild-type cells at a ratio of 35:1 and preincubated with 500 nM MβCD for 2 h and then incubated with Dox for 24 h. (**e, f**) Type I and II harbouring Src-EGFP were grown in the presence of Dox for indicated time periods. Lysates from these cells were subjected to the DRM separation experiment to separate DRM and non-DRM fractions. The fractions were analysed by immunoblotting using indicated antibodies. Flotillin1 and caveolin1 were used as DRM marker, and transferrin receptor (TfR) was used as a non-DRM marker. Ratio of DRM localisation of Src-EGFP was calculated by DRM fraction / total amount. The mean ratios ± SDs were obtained from three independent experiments. *, *P* < 0.05; **, *P* < 0.01; ***, *P* < 0.001; NS, not significantly different; two-way ANOVA was calculated compared with the Dox-treated cells (a) or non-treated cells (c, f). The scale bars indicate 50 μm.

To verify this possibility, we forcibly localised Src in either lipid rafts or non-lipid rafts using a rapamycin-inducible FKBP-FRB dimerisation system (**Fig. 3a and Supplementary Fig. 3a**). Before analysis, we confirmed that rapamycin treatment did not affect Src-induced delamination (**Supplementary Fig. 3b, c**). DRM separation analysis showed that rapamycin treatment induced Src translocation into lipid rafts in cells expressing Pag1TM-FKBP, but not Pag1TM^mut^-FKBP (**Fig. 3b**). In a two-dimensional cell competition assay, Src trapped in lipid rafts promoted basal delamination (**Fig. 3c, d and Supplementary Fig. 3d**), in a manner similar to Src-EGFP overexpression. In contrast, Src trapped outside lipid rafts induced apical extrusion (**Fig. 3c, d and Supplementary Fig. 3d**). Similar phenomena were observed in the three-dimensional culture of mosaic cysts (**Fig. 3e, f**). Importantly, when this system was introduced in MDCK type II cells, forced localisation of Src into lipid rafts conferred the ability of basal delamination to Src-expressing cells (**Supplementary Fig. 3e**). These data support our hypothesis that lipid raft localisation of activated Src is crucial for inducing basal delamination.

**Figure 3.**
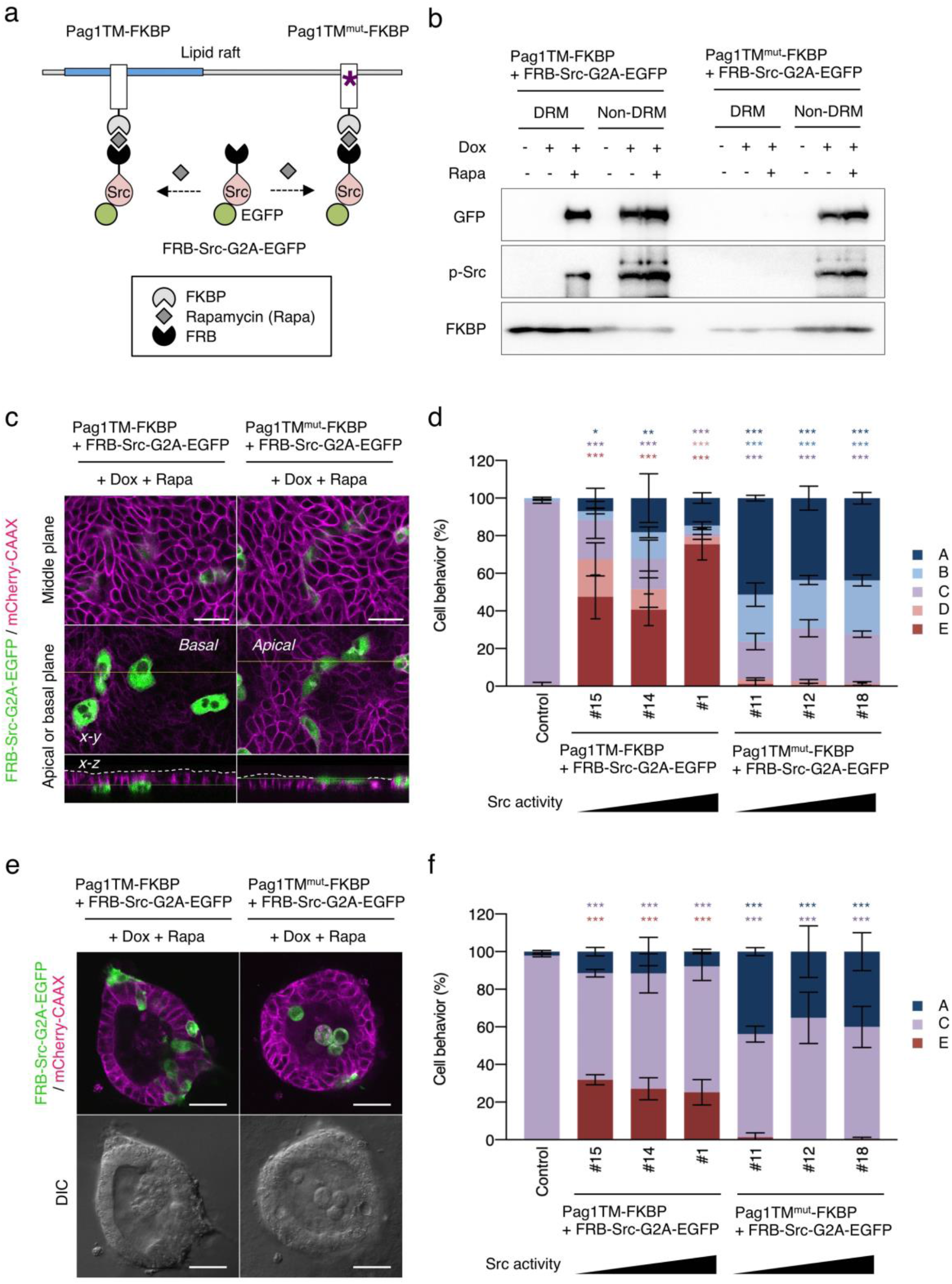
Lipid raft localisation of Src is a critical determinant for basal delamination. (**a**) Schematic model of Src recruiting system using rapamycin-mediated FKBP-FRB dimerization. An asterisk indicates point mutations of palmytoilation sites. (**b**) MDCK type I cells harbouring FRB-Src-G2A-EGFP and Pag1TM-FKBP or Pag1TM^mut^-FKBP were incubated in the presence or absence of 1 μg/ml Dox and 100 nM rapamycin (Rapa) for 24 h. Lysates from these cells were subjected to the DRM separation experiment to separate DRM and non-DRM fractions. The fractions were analysed by immunoblotting using indicated antibodies. (**c, d**) Type I were mixed with wild-type cells at a ratio of 35:1 and incubated in the presence of Dox and Rapa for 24 h. Cell behaviour was assessed by using the criteria (Fig. 1c). (**e, f**) Type I cells harbouring FRB-Src-G2A-EGFP and Pag1TM-FKBP or Pag1TM^mut^-FKBP were mixed with wild-type cells at a ratio of 8:1, the mosaic cysts were grown in collagen matrix in the presence of Dox and Rapa for 2 days. Cell behaviour was assessed by three categories. The mean ratios ± SDs were obtained from three independent experiments. *, *P* < 0.05; **, *P* < 0.01; ***, *P* < 0.001; two-way ANOVA was calculated compared with the non-treated cells. The scale bars indicate 50 μm.

### The STAT3-MMP axis is crucial for Src-induced basal delamination

To determine the downstream pathways of Src activation in lipid rafts, we performed an inhibitor screening assay using Src-EGFP-expressing MDCK type I cells. In the cell competition assay, we found that pretreatment with specific STAT3 inhibitors significantly suppressed Src-induced basal delamination (**Fig. 4a**). The same result was obtained when dominant-negative STAT3 (STAT3-DN) was overexpressed (**Fig. 4b**). STAT3 was activated (p-STAT3, pTyr705) by Src expression, but not by Ras (**Fig. 1a**), and this activation was elevated under competitive conditions (**Fig. 4c, d**). Comparative analysis of MDCK type I and II cells showed that STAT3 was markedly activated by Src expression in MDCK type I cells, whereas STAT3 activation was significantly lower in type II cells than in type I cells, even though intracellular tyrosine phosphorylation levels (pY1000 blot) were almost comparable (**Fig. 4e, f**). These results suggest that selective STAT3 activation via Src in lipid rafts is tightly linked with Src-induced basal delamination.

**Figure 4.**
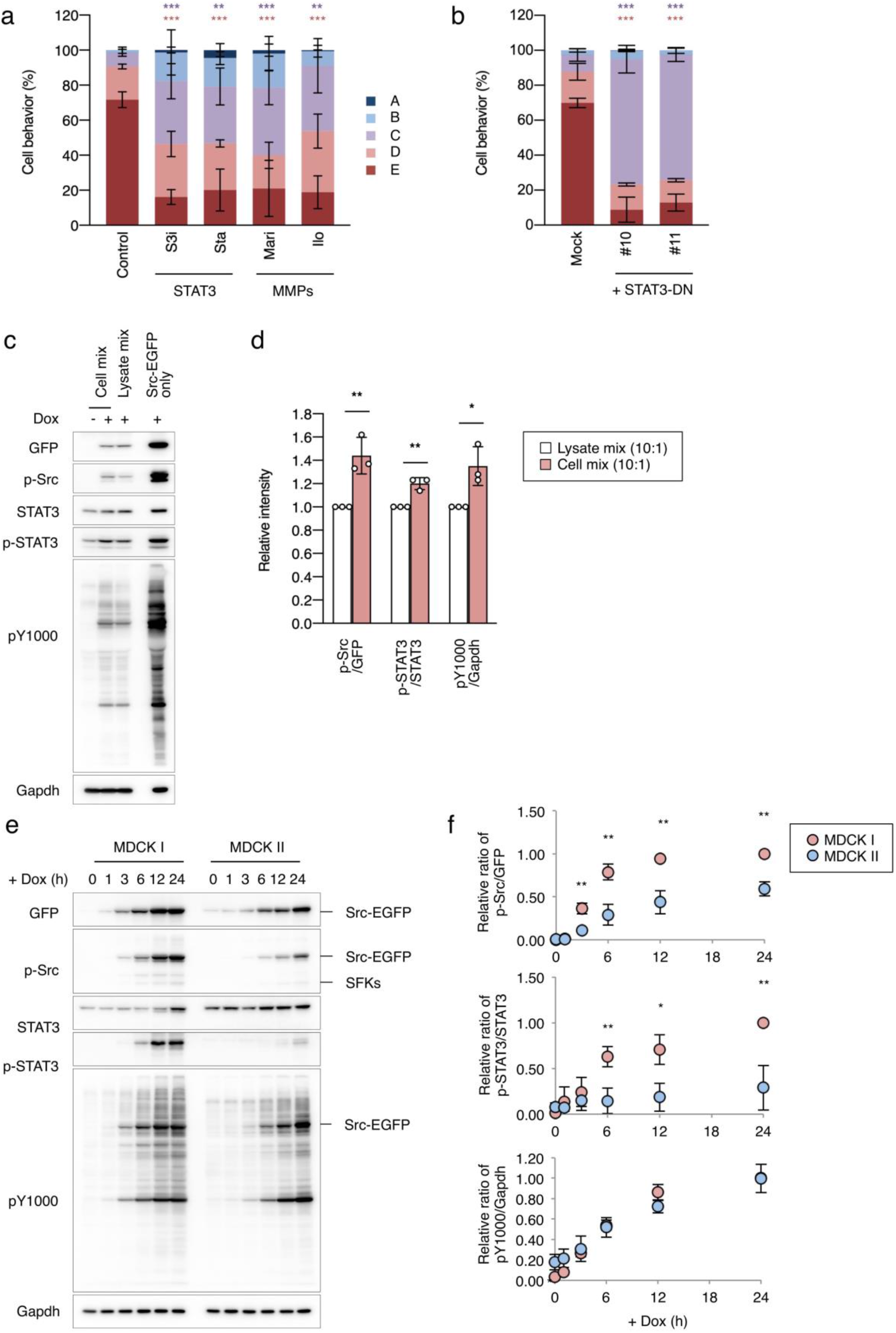
Src induces basal delamination through activation of the STAT3-MMP axis. (**a**) MDCK type I cells harbouring Src-EGFP were mixed with wild-type cells at a ratio of 35:1 and preincubated with the 100 μM S3i-201 (S3i), 20 μM stattic (Sta), 20 μM marimastat (Mari), or 400 nM ilomastat (Ilo) for 2 h and then incubated with Dox for 24 h. (**b**) Two type I clones harbouring Src-EGFP and STAT3-Y705F (DN) were mixed with wild-type cells at a ratio of 35:1 and incubated with Dox for 24 h. Cell behaviour was assessed by using the criteria (Fig. 1c). (**c, d**) Type I cells harbouring Src-EGFP were mixed with wild-type cells at a ratio of 10:1 and incubated with Dox for 24 h. Wild-type and Src-EGFP harbouring cells were separately incubated with Dox for 24 h, and cell lysates were mix at a ratio of 10:1. These cell lysates were subjected to immunoblotting analysis using the indicated antibodies. Relative intensity of p-Src (pY418)/GFP, p-STAT3 (pY705)/STAT3, and pY1000/Gapdh were calculated by setting the value for lysate mix to one. (**e, f**) Type I and II harbouring Src-EGFP were incubated in the presence of Dox for indicated time periods. Lysate from these cells were analysed by immunoblotting using indicated antibodies. Relative ratio of p-Src/GFP, p-STAT3/STAT3, and pY1000/Gapdh were calculated by setting the value for 24 h-treated type I cells to one. The mean ratios ± SDs were obtained from three independent experiments. *, *P* < 0.05; **, *P* < 0.01; ***, *P* < 0.001; two-way ANOVA was calculated compared with the Dox-treated cells (a) or mock-transfected cells (b), and two-tailed *t*-test was calculated (d, f).

STAT3 activated by lipid raft Src has been reported to upregulates MMP expressions^26^. Accordingly, we analysed the contribution of MMPs to Src-induced basal delamination. Src-induced basal delamination was suppressed by pretreatment with broad-spectrum MMP inhibitors (**Fig. 4a**), indicating that the Src-transformed cells delaminate into basement membrane through MMP-mediated collagen matrix degradation (**Supplementary Fig. 2e**). We then investigated the expression of MMPs in MDCK type I and II cells. After Src induction, a broad range of MMPs including both secreted-(MMP3, MMP11) and membrane-type (MMP14 and MMP17), were significantly upregulated in MDCK type I cells, whereas the levels of MMP upregulation were substantially lower in MDCK type II cells (**Supplementary Fig. 4a**). Further, upregulation of MMP14 and MMP17 was suppressed by pretreatment with the STAT3 inhibitor, S3i-201 (**Supplementary Fig. 4b**), indicating that the expression of MMP genes is upregulated by STAT3 activation. Immunofluorescence also showed that MT4-MMP, a gene product of MMP17, was upregulated and accumulated in the focal adhesions of Src-expressing cells in a two-dimensional cell competition assay (**Supplementary Fig. 4c**). These observations demonstrate that Src activation in lipid rafts induces STAT3 activation, followed by upregulation of MMPs required for basal delamination.

### CDCP1 induces basal delamination by recruiting Src in lipid rafts

We next considered how Src is selectively localised to lipid rafts in MDCK type I cells. In the two MDCK cell types, there was no significant difference in the cholesterol contents (**Fig. 2c**) and expression levels of lipid raft marker proteins, flotillin and caveolin, in the DRM fractions (**Fig. 2e and Supplementary Table 2**). These results indicate that lipid rafts structures are seemingly comparable between these two cell types, suggesting that differences in the lipid raft Src localisation may arise due to differences in scaffold proteins that recruit Src to the lipid rafts. To identify the Src scaffold protein(s) in lipid rafts, we isolated DRM fractions from Src-inducible MDCK type I and II cells and analysed their protein contents by mass spectrometry (**Supplementary Fig. 5a**). After Src overexpression, the accumulated protein in DRM fractions was higher in MDCK type I cells than in type II cells (**Supplementary Fig. 5a and Supplementary Table 3**). Among 105 proteins enriched in the DRM fractions from MDCK type I cells, we chose CDCP1 and Met as potential Src scaffold protein candidates in lipid rafts (**Supplementary Fig. 5a and Supplementary Table 3**) and analysed their functions.

Previous studies have already shown that CDCP1 is associated with activated Src in lipid rafts (**Fig. 5a**)^26^. Under cell competition conditions, we found that CDCP1 accumulated in the plasma membrane of Src-expressing cells, suggesting that CDCP1 contributes to Src regulation in the context of cell competition (**Supplementary Fig. 5b**). We thus examined the role of CDCP1 in CDCP1-inducible MDCK type I cells. In the cell competition assay, overexpression of wild-type CDCP1 induced a morphological change to a stretched shape and delamination into the basement membrane (**Fig. 5a-c**). Overexpression of the CDCP1-Y734F mutant (CDCP1-YF), which has a mutation in the Src-association site, showed no obvious effects (**Fig. 5a-c**). In contrast, overexpression of the CDCP1-C689G-C690G mutant (CDCP1-CG), which has mutations in the palmitoylation sites required for lipid raft localisation, induced apical extrusion (**Fig. 5a-c**). Similar phenomena were observed in the three-dimensional cell competition assay (**Fig. 5d, e**). These data demonstrate that CDCP1 functions as an activating scaffold for endogenous Src in lipid rafts and support our model that lipid raft localisation of activated Src induces basal delamination instead of apical extrusion.

**Figure 5.**
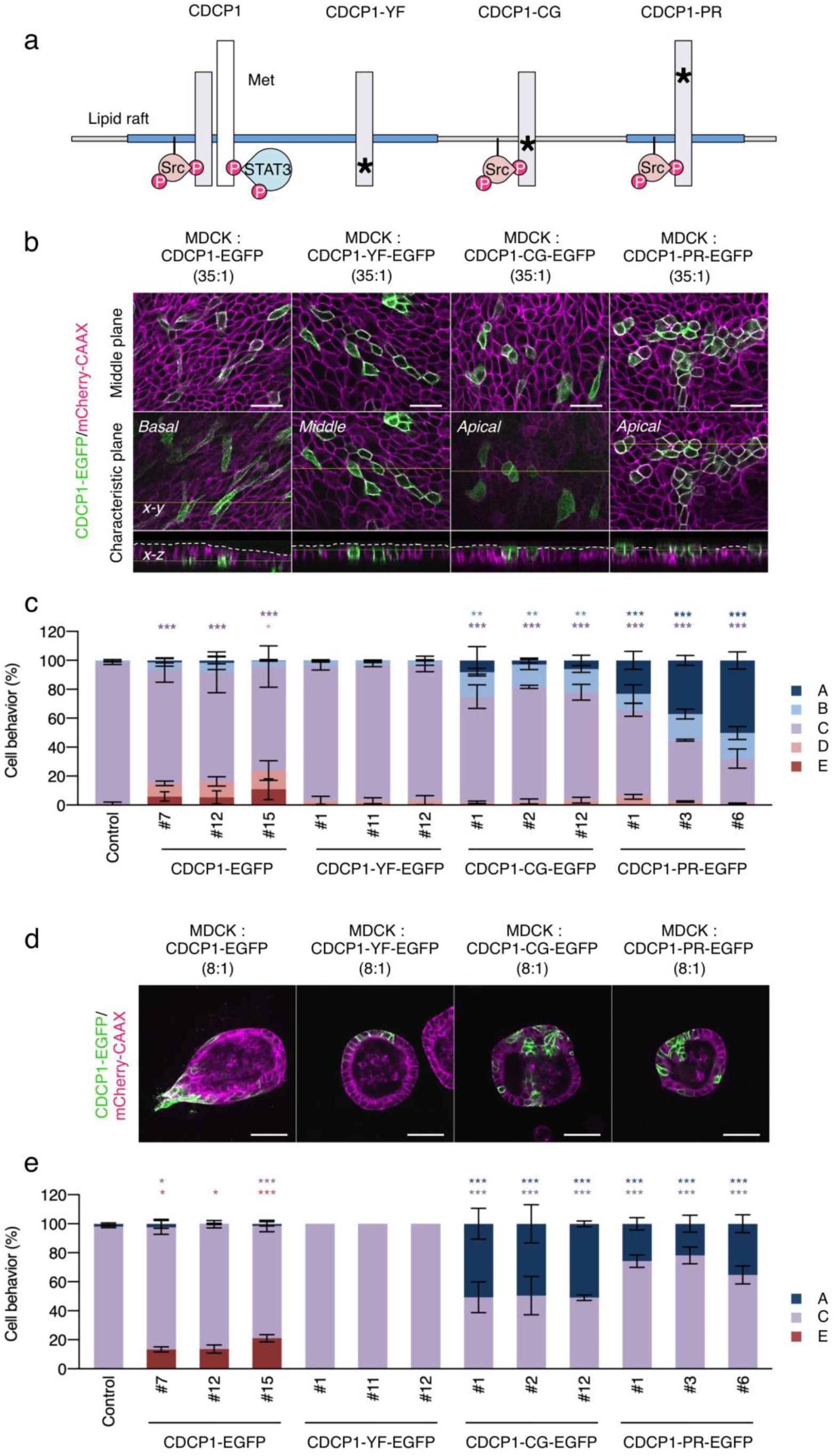
CDCP1 induces basal delamination through Src activation within lipid rafts. (**a**) Schematic model of CDCP1 and its mutants. Blue squares indicate lipid raft domains, and asterisks indicate point mutations. (**b, c**) MDCK type I cells harbouring CDCP1-EGFP, CDCP1-YF-EGFP, CDCP1-CG-EGFP, or CDCP1-PR-EGFP were mixed with wild-type cells at a ratio of 35:1 and incubated with Dox for 24 h. Cell behaviour was assessed by using the criteria (Fig. 1c). (**d, e**) Type I cells harbouring CDCP1-EGFP, CDCP1-YF-EGFP, CDCP1-CG-EGFP or CDCP1-PR-EGFP were mixed with wild-type cells at a ratio of 8:1, the mosaic cysts were grown in collagen matrix in the presence of Dox for 2 days. Cell behaviour was assessed by three categories. The mean ratios ± SDs were obtained from three independent experiments. *, *P* < 0.05; **, *P* < 0.01; ***, *P* < 0.001; two-way ANOVA was calculated compared to the control cells (c) or mosaic cysts (e). The scale bars indicate 50 μm.

On the contrary, although Met inhibitors suppressed Src-induced basal delamination (**Supplementary Fig. 5c**), overexpression of Met alone did not induce endogenous Src activation or basal delamination (**Supplementary Fig. 5d-f**), indicating that Met activity is required but not sufficient for cell delamination. The lipid raft-localised CDCP1-Src complex associates with Met for activating the STAT3 pathway, and an efficient interaction of CDCP1-Met is known to requires proteolytic shedding of CDCP1 extracellular domain^26^. To examine whether the CDCP1-Met-STAT3 axis is functional even in the context of cell competition, we introduced K365A-R368A-K369A mutations in CDCP1 (CDCP1-PR) to repress proteolytic shedding and interaction with Met and STAT3 activation (**Fig. 5a**). In both two- and three-dimensional cell competition, overexpression of CDCP1-PR induced apical extrusion (**Fig. 5b-e**). Further, CDCP1-induced basal delamination was significantly suppressed by pretreatment with specific inhibitors of Met, STAT3, and MMPs, and a part of the CDCP1-overexpressing cells were extruded into the apical side in two- and three-dimensional cell competition assays (**Supplementary Fig. 5g, h**). These data suggest that Src-induced basal delamination is spatially controlled by CDCP1 through association with Met, followed by activation of the STAT3-MMPs axis via lipid rafts (**Supplementary Fig. 5i**).

### Activation of the CDCP1-STAT3-MMPs axis induces invasion ability in renal cancer cells

In the three-dimensional cell competition assay, CDCP1-overexpressing MDCK type I cells protruded from the cell layer of cysts and migrated into the ECM along with normal cells (**Fig. 5d**). As these phenotypes appear quite similar to cancer invasion, we assessed the role of the CDCP1-Src complex in cancer cells using spheroid cultures of the renal cancer cell line A498, in which CDCP1 was highly expressed and was further upregulated by HGF treatment (**Supplementary Fig. 6a-c**). CDCP1 knockdown attenuated Met expression and partially suppressed the activation of HGF signalling components, including Src family kinases (SFKs), STAT3, and MMP expression (**Supplementary Fig. 6a, b, d**). Furthermore, CDCP1 knockdown suppressed HGF-induced morphological changes accompanied with N-cadherin upregulation, a marker of epithelial-mesenchymal transition (EMT), resulting in the inhibition of *in vitro* invasion activity (**Supplementary Fig. 6c, e**). These data suggest that CDCP1 supports activation of HGF-Met signalling, which induces EMT-mediated invasive phenotypes by activating the Src-STAT3-MMPs axis, even in cancer cells. To further elucidate the role of CDCP1 upregulation in heterogeneous cancer cell populations, parental A498 (CDCP1^High^; CMFDA-stained) cells were mixed with CDCP1-knockdown A498 (CDCP1^Low^; unstained) cells, and the mosaic spheroids were cultured in a collagen matrix (**Fig. 6a**). After HGF stimulation, CDCP1^High^ cells, but not CDCP1^Low^ cells, protruded from the spheroids and migrated into the collagen matrix (**Fig.6b, c**). Furthermore, some CDCP1^High^ cells co-migrated together with a mass of CDCP1^Low^ cells (**Fig. 6b**), implying that highly invasive CDCP1^High^ cells function as tip cells and lead collective cell invasion. To confirm the molecular mechanism downstream of CDCP1, we re-expressed CDCP1 or its mutants in CDCP1^Low^ cells. The defect in HGF-induced invasion was recovered by re-expression of wild-type CDCP1, but not the CDCP1-YF mutant, and was partially recovered by re-expression of CDCP1-CG mutants (**Fig. 6d, e**). Additionally, the HGF-induced invasive phenotype of CDCP1^High^ cells was suppressed by pretreatment with inhibitors of SFKs, STAT3, MMPs, and cholesterol synthesis (**Fig. 6f, g**). These observations suggest that upregulated CDCP1 promotes HGF-induced collective invasion of cancer cells via activation of the Src-STAT3-MMPs axis through the lipid raft signalling platform.

**Figure 6.**
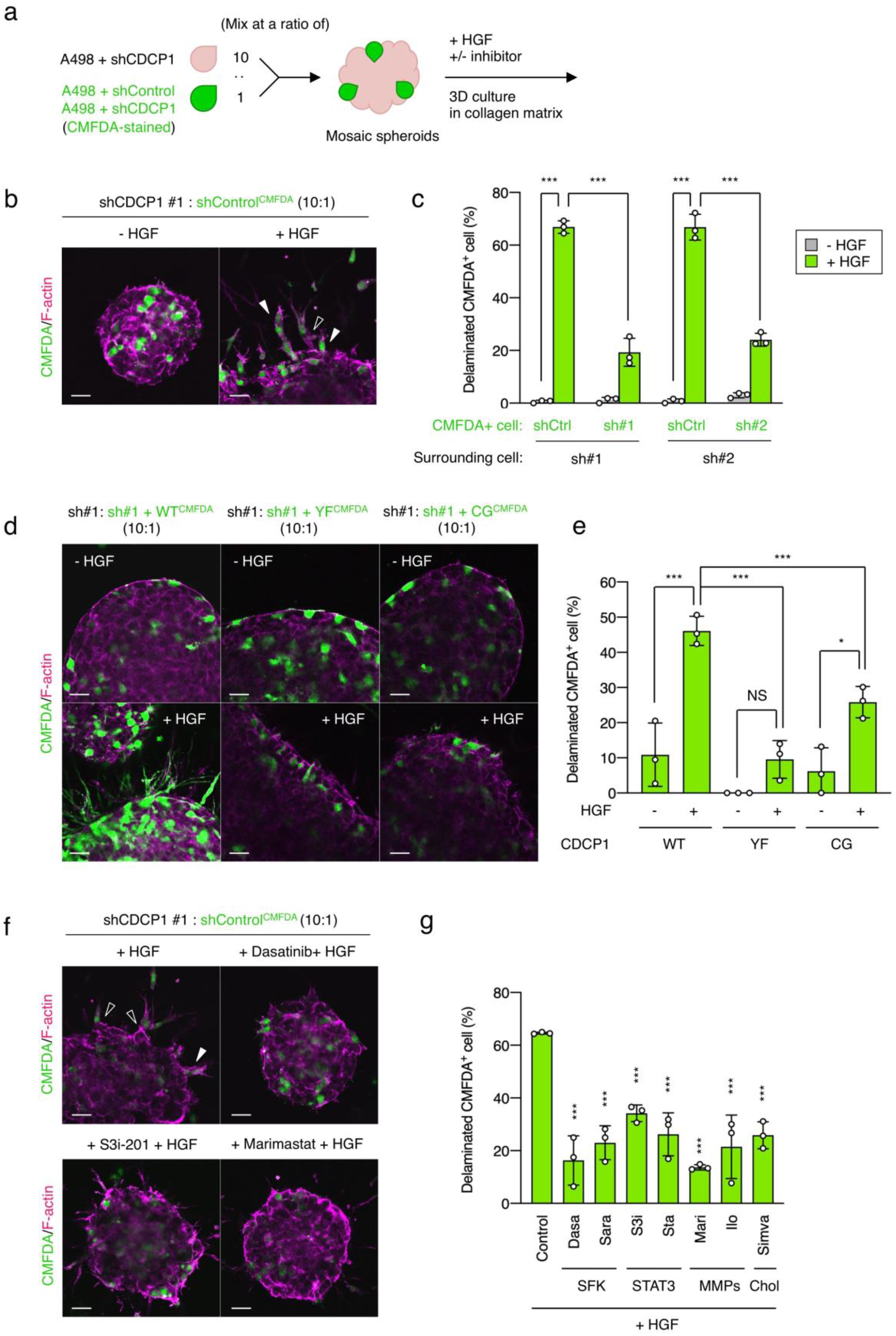
CDCP1-expressing cancer cells lead collective invasion with low-invasive cells. (**a**) Schematic diagram of mosaic spheroid analysis of A498 cells. shControl or shCDCP1 expressing A498 cells were stained using CMFDA and mixed with non-stained shCDCP1 expressing A498 cells at a ratio of 1:10, and the mosaic spheroids were embedded within collagen matrix and grown in the presence of HGF and/or inhibitors. (**b, c**) A498 mosaic spheroids embedded within collagen matrix were incubated in the presence or absence of 100 ng/ml HGF for 2 days. After fixation, filamentous actin was stained with Alexa594-conjugated phalloidin. Percentage of delaminated CMFDA-stained A498 cells was calculated by delaminated cell / total peripherally localised cells (n > 60). The mean ratios ± SDs were obtained from three independent experiments. (**d, e**) A498-shCDCP1 cells harbouring CDCP1, CDCP1-YF, or CDCP1-CG were stained using CMFDA and mixed with non-stained A498-shCDCP1 cells at a ratio of 1:10, and the mosaic spheroids were embedded within collagen matrix and grown in the presence of HGF. The mean ratios ± SDs were obtained from three independent experiments. (**f, g**) A498 mosaic spheroids were pretreated with 10 nM dasatinib (Dasa), 20 μM saracatinib (Sara), 100 μM S3i-201 (S3i), 20 μM stattic (Sta), 20 μM marimastat (Mari), 400 nM ilomastat (Ilo) or 1.6 μM simvastatin (Simva) for 2 h and incubated in the presence of 100 ng/ml HGF for 2 days. The mean ratios ± SDs were obtained from three independent experiments. *, *P* < 0.05; ***, *P* < 0.001; NS, not significantly different; ANOVA was calculated compared to the HGF-treated mosaic spheroids. The scale bars indicate 50 μm.

## Discussion

In this study, we demonstrated that spatial regulation of Src within the plasma membrane is crucial for cell fate decisions under cell competition status. In the cell competition assay, Src-transformed MDCK type II cells were apically extruded, whereas Src-transformed type I cells underwent basal delamination. Comparative analysis of these two cell types revealed that Src localisation in lipid rafts is a critical determinant of basal delamination through activation of the STAT3-MMP axis and maturation of focal adhesions (**Supplementary Fig. 5j**). Furthermore, we showed that lipid raft localisation of Src is mediated by the transmembrane scaffold, CDCP1, and that cancer cells with CDCP1-Src upregulation behave as leader cells to promote collective cancer invasion in heterogeneous populations. These findings suggest that upregulation of Src activity via lipid raft scaffolds such as CDCP1, may facilitate escape from apical extrusions by inducing invasive potential, leading to promotion of cancer malignancy.

As Src has a strong oncogenic potential, Src protein expression and/or activity is strictly regulated via phosphorylation by Csk^29^, the ubiquitin-proteasome system, and exosomal exclusion^30^. Additionally, intracellular distribution of Src contributes to regulate its function. Src is anchored to the membrane via myristoylation. Upon activation, part of Src is translocated into lipid rafts via scaffold proteins, such as the negative regulator Pag1/Cbp^22, 25^ and positive regulator CDCP1^28^, and is inactivated or activated on scaffold proteins, depending on the cellular context. In this study, we found that activated Src efficiently accumulated in lipid rafts to activate the STAT3 pathway in MDCK type I cells, whereas Src accumulation in lipid rafts was substantially attenuated in MDCK type II cells. This difference was tightly associated with differential phenotypes in cell competition, basal delamination in type I cells, and apical extrusion in type II cells. There are two potential explanations for this difference. First, differential phenotypes may arise due to differences in the compositions of lipid rafts, consisting mainly of cholesterol and sphingolipids. Although there were no significant differences in cholesterol content between the two cell lines, differences in sphingolipid content may influence lipid raft integrity and retention of activated Src^23^. Indeed, differences in sphingolipid species between the two cell lines have been reported previously^31^. Furthermore, sphingolipids and cholesterol metabolism is disordered in various cancers and is associated with the promotion of cancer malignancy^32-34^. In a cell competition mouse model, apical extrusion of Ras-transformed cells from epithelial tissues was suppressed by obesity^8^. These lines of evidence suggest that alterations in lipid metabolism may also allow abnormal/mutated cells to escape apical extrusion in the initial phase of carcinogenesis.

Another potential cause for inducing differential phenotypes between MDCK type I and II cells is the differential content of Src scaffolds in lipid rafts. In this study, we revealed that CDCP1 plays an important role as an Src scaffold in lipid rafts to control cell fate in the context of cell competition. Overexpressed CDCP1 recruited activates Src into lipid rafts, followed by activation of the STAT3-MMP axis through association with Met^26^. However, Src relocation was not completely suppressed by CDCP1 knockout (data not shown), suggesting that an additional Src scaffold(s) exists in lipid rafts. Among the DRM-enriched proteins identified from MDCK type I cells, EphA2 (**Supplementary Table 3**), a receptor tyrosine kinase that is associated with Src^35^ and contributes to cancer cell invasion^36^, is another strong candidate. In a cell competition assay with Ras-transformed cells, EphA2 is involved in regulating Src and its downstream signalling^37^. These observations imply that EphA2 may function cooperatively with CDCP1 to induce basal delamination in MDCK type I cells.

Upregulation of CDCP1 is frequently observed in accordance with cancer development^26, 27, 38^. CDCP1 was highly expressed in patients with renal cancer in The Cancer Genome Atlas (TCGA) cohort (**Supplementary Fig. 7a**), and high expression of CDCP1 was correlated with poor prognosis (**Supplementary Fig. 7b**). The STAT3 pathway is also activated in various cancers to promote cancer malignancy^39^. In this study, we further found that expression of MMP14/MT1-MMP and MMP17/MT4-MMP was increased by STAT3 activation in MDCK cells. Both MMPs are expressed in A498 renal cancer cells and may be involved in cancer metastasis. MMP14 is involved in cell extrusion of transformed cells from the normal epithelial layer^40^. MMP17 is a GPI-anchored membrane-type MMP involved in cell invasion^41, 42^ and is upregulated in various cancers including breast, gastric, and colon cancer^43, 44^. MMP17 was also highly expressed in renal cancer in the TCGA dataset, and its expression was elevated during cancer progression (**Supplementary Fig. 7c**). Furthermore, high MMP17 expression was associated with poor prognosis in patients with renal cancer (**Supplementary Fig. 7d**). These data suggest that MMP17 is an important factor in the malignant progression of renal cancer.

In summary, comparative analysis of contrasting phenotypes in Src-activated MDCK cells revealed that activated Src in lipid rafts facilitates cellular escape apical extrusions by inducing invasive potential in epithelial cells. Further analysis in cancer cells also showed that Src activation by upregulation of lipid raft scaffolds promoted invasive activity during cancer development. Based on these findings, we propose that the regulatory system of Src distribution in lipid rafts, such as raft scaffold proteins and lipid contents, may provide promising therapeutic targets to control carcinogenesis and cancer progression.

## Materials and methods

### Cell and cell culture

Two types of MDCK and A498 cells were cultured in Dulbecco’s modified Eagle’s medium (DMEM) supplemented with 10% fetal bovine serum (FBS) at 37°C in a 5% CO_2_ atmosphere. MDCK type II cells were gifted from Dr. Fujita^12^. Recombinant HGF (100-39) was purchased from PeptroTech.

### Plasmid construction and gene transfer

Inducible gene expression system based on Tet-On 3G (Clontech) was employed. Mouse Src and human CDCP1 were fused at 5’ terminus of EGFP (or EBFP) and introduced into pRetroX-TRE3G plasmid. Src-KD was generated by point mutation of K297M by mutagenesis PCR. CDCP1 mutants were generated by point mutations of Y734F, C689G-C690G, or K365A-R368A-K369A by mutagenesis PCR. Human Met and HRASV12 were introduced into pRetroX-TRE3G plasmid. For construction of plasmids for FKBP-FRB dimerization (**Supplementary Figure 3a**), transmembrane domain (63 residues of N-terminus) of mouse Pag1/Cbp was amplified, and point mutations (C39G-C42G; Pag1TM^mut^) were generated by mutagenesis PCR. The Pag1TM or Pag1TM^mut^ DNA fragments were fused to 5’ terminus of human FKBP12 and introduced into pCX4 plasmid. FRB domain (2021-2113 residues) of human MTOR was amplified, and the FRB domain was fused with 5’ terminus of cytosolic Src-EGFP (Src-G2A-EGFP) and introduced into pRetroX-TRE3G plasmid. To construct mCherry-CAAX plasmid, CAAX motif (21 residues of C-terminus) of human KRAS was amplified, and the DNA fragment was fused to 3’ terminus of mCherry and introduced into pCX4 plasmid. FAK-DN and STAT3-DN were generated by point mutation of Y397F and Y705F, respectively, by mutagenesis PCR. Construction of pCX4-STAT3-MER plasmid was described in previous report^26^. Gene transfer of pRetroX-TRE3G and pCX4 was carried out by retroviral infection. Retroviral production and infection were performed as described previously^45^. shRNAs against human CDCP1 (TRCN0000134829 and TRCN0000137203, Sigma-Aldrich) and MMP17 (TRCN0000049976 and TRCN0000049977, Sigma-Aldrich) were introduced in to cells through lentiviral infection using packaging mix (Sigma-Aldrich).

### Cell competition analysis

For two-dimensional competition assay, wild-type and gene expression inducible system harbouring MDCK cells were mixed at a ratio of 35:1 and spread on collagen sheet using Cellmatrix type I-A (Nitta Gelatin) in DMEM supplemented with 10% FBS. Cell Tracker Green CMFDA (C7025, Thermo Fisher Scientific) was used for labelling before expression of fluorescent protein. For collagen degradation assay, a collagen sheet containing 2% DQ-collagen type I (D12060, Thermo Fisher Scientific) was used. After MDCK sheet formation, cells were incubated in DMEM supplemented with 5% FBS and 1 μg/ml doxycycline (Dox). For three-dimensional competition assay, wild-type and gene expression inducible system harbouring cells were mixed at a ratio of 8:1 and cultured in low-attachment dish (Sumitomo Bakelite) for 2 days. Mosaic spheroids were harvested and recultured on Matrigel (356231, Corning) for 3 days. After cystogenesis, collagen type I matrix was overlaid and incubated in DMEM supplemented with 5% FBS and 1 μg/ml Dox for 2 days.

### Mosaic spheroid analysis

A498 cells harbouring shControl were labelled with CMFDA and mixed with A498 cells harbouring shCDCP1 at a ratio of 1:10 (**Figure 6a**). Mixed cells were cultured in low cell adhesion dish for 2 days. Mosaic spheroids were harvested and recultured in collagen type I matrix in the presence of 50 ng/ml HGF for 2 days.

### Immunoblotting analysis

Cell were lysed in n-octyl-b-D-glucoside (ODG) buffer [20 mM Tris-HCl (pH7.4), 150 mM NaCl, 1 mM EDTA, 1 mM Na3VO4, 20 mM NaF, 1% Nonidet P-40, 5% glycerol, 2% ODG and protease inhibitor cocktail (Nacalai Tesque)], and immunoblotting was performed using primary antibodies listed below. All blots were visualized and quantified using a LuminoGraph II (Atto).

### Immunofluorescent microscopic analysis

MDCK cells, cysts and A498 spheroids were fixed with 4% paraformaldehyde (PFA) and blocked with 1% BSA. Fixed MDCK cells were incubated with primary antibodies listed below and then incubated further with Alexa Fluor 488-conjugated antibody (Thermo Fisher Scientific). Fixed A498 cells were incubated with Alexa Fluor 594-conjugated phalloidin (Thermo Fisher Scientific). Immunostained cells were observed using a FV1000 confocal fluorescence microscope system (Olympus).

### Antibodies and inhibitors

The following primary antibodies were used in this study: anti-STAT3 (9132), anti-STAT3 pY705 (9145), anti-phospho-tyrosine pY1000 (8954), anti-Met (3127, clone 25H2) and anti-Met pY1234/1235 (3077) were purchased from Cell Signaling Technologies. Anti-GAPDH (sc-32233, clone 6C5), anti-FAK pY576/577 (sc-21831), anti-SFK (sc-18, clone SRC2), and anti-EPLIN (sc-136399, clone 20) were purchased from Santa Cruz Biotechnology. Anti-flotillin1 (610821), anti-caveolin1 (610407), and anti-paxillin (610051) were purchased from BD bioscience. Anti-Ras (OP01, clone Y13-259) and anti-cortactin (05-180, clone 4F11) were purchased from Millipore. Anti-Src pY418 (44-655G) and anti-GFP (A6455) were purchased from Thermo Fisher Scientific. Anti-FKBP12 (ab2918) and anti-MT4-MMP (ab51075) were purchased from abcam. Anti-CDCP1 (LS-C172540, clone 5B3) was purchased from LSBio. Anti-HIF-1α (NB100-105) was purchased from Novas Biologicals.

The following inhibitors were used in this study: Fumonisin B1 (344850), S3i-201 (573102), stattic (573099), and AMG-1 (448104) were purchased from Calbiochem. FAK inhibitor 14 (SML0837), marimastat (M2699), myriocin (M1177), and simvastatin (S6196) were purchased from Sigma-Aldrich. D-threo-1-phenyl-2-decanoylamino-3-morpholino-1-propanol (1756) was purchased from Matreya. Lovastatin (438185) was purchased from Millipore. Dasatinib (BMS-354825, 1586-100) was purchased from BioVision. Saracatinib (AZD0530, S1006) was purchased from Selleck. Rapamycin (30037-94) was purchased from Nacalai Tesque. Other inhibitors were obtained from the Screening Committee of Anticancer Drugs.

### DRM separation analysis

Cells were lysed with homogenization buffer [50 mM Tris-HCl (pH7.4), 150 mM NaCl, 1 mM EDTA, 1 mM Na3VO4, 20 mM NaF, 0.25% Triton X-100 and protease inhibitor cocktail (Nacalai Tesque)] and separated on a discontinuous sucrose gradient (5-35-40%) by ultracentrifugation at 150,000 *g* using Optima L-100XP (Bechman Coulter). A total of 11 fractions were collected from the top of the sucrose gradient.

### Cholesterol analysis

Cholesterol amount was analysed using the Amplex Red Cholesterol Assay Kit (A12216, Thermo Fisher Scientific) according to the manufacturer’s protocol. The assay was performed to cell lysates from DRM fractions that were normalized for protein concentration.

### Quantitative real-time PCR analysis

Total RNA from cells was prepared using Sepasol-RNA I Super G (Nacalai Tesque) and cDNA was prepared with the ReverTra Ace qPCR RT Master Mix (FSQ-201, Toyobo). Real-time PCR was performed on a QuatStudio 5 (Thermo Fisher Scientific) using Thunderbird SYBR qPCR Mix (QPS-201, Toyobo). The expression of the housekeeping gene GAPDH was used to normalized the amount of total RNA. The primers used in this study are listed in Supplementary Table 4.

### Clinical and gene expression analysis

Clinical and RNA-seq data of renal clear cell carcinoma (606 patients) from the TCGA dataset were used. Survival curves were constructed using Kaplan-Meier method and compared by using the log-rank test.

### In vitro invasion assay

Matrigel invasion chambers (354480, Corning) were used for the invasion assays. 1 × 10^5^ cells were seeded on chambers containing culture media with or without 100 ng/ml HGF. After incubation at 37°C for 24 h, invaded cells were fixed with 4% PFA and then stained with 1% crystal violet. Invaded cells were counted on micrographs; in each experiment, cells were counted on five randomly chosen fields.

### Statistics and reproducibility

For data analysis, unpaired two-tailed *t*-test were performed to determine the *P*-values. For multiple group comparisons, two-way analysis of variance (ANOVA) was used. A *P*-value of < 0.05 was considered to reflect a statistically significant difference. All data and statistics were derived from at least three independent experiments.

## Acknowledgments

We thank Dr. Kajita for experimental support and Dr. Saito for mass spectrometry analysis. The inhibitor kit was provided by the Screening Committee of Anticancer Drugs, which is supported by Grant-in-Aid for Scientific Research on Innovative Areas, Scientific Support Programs for Cancer Research, from The Ministry of Education, Culture, Sports, Science and Technology of Japan. This work was supported by a Grant-in-Aid for Scientific Research (C) (19K07639, to K.K.) and Grant-in-Aid for Scientific Research on Innovative Areas (26114001, to M.O.) from The Japan Society for the Promotion of Science.

## Competing interests

The authors declare that they have no conflict of interest.

## Author contributions

K.K. designed study and performed most experiments and data analysis. P.C. conducted experiments using CDCP1. S.K and Y.F. supported experiments. K.K. and M.O. wrote the manuscript with input from all authors.

## Notes

### Competing Interest Statement

The authors have declared no competing interest.

