## Supplemental Figures for "Src activation in lipid rafts confers epithelial cells with invasive potential to escape from apical extrusion during cell competition"

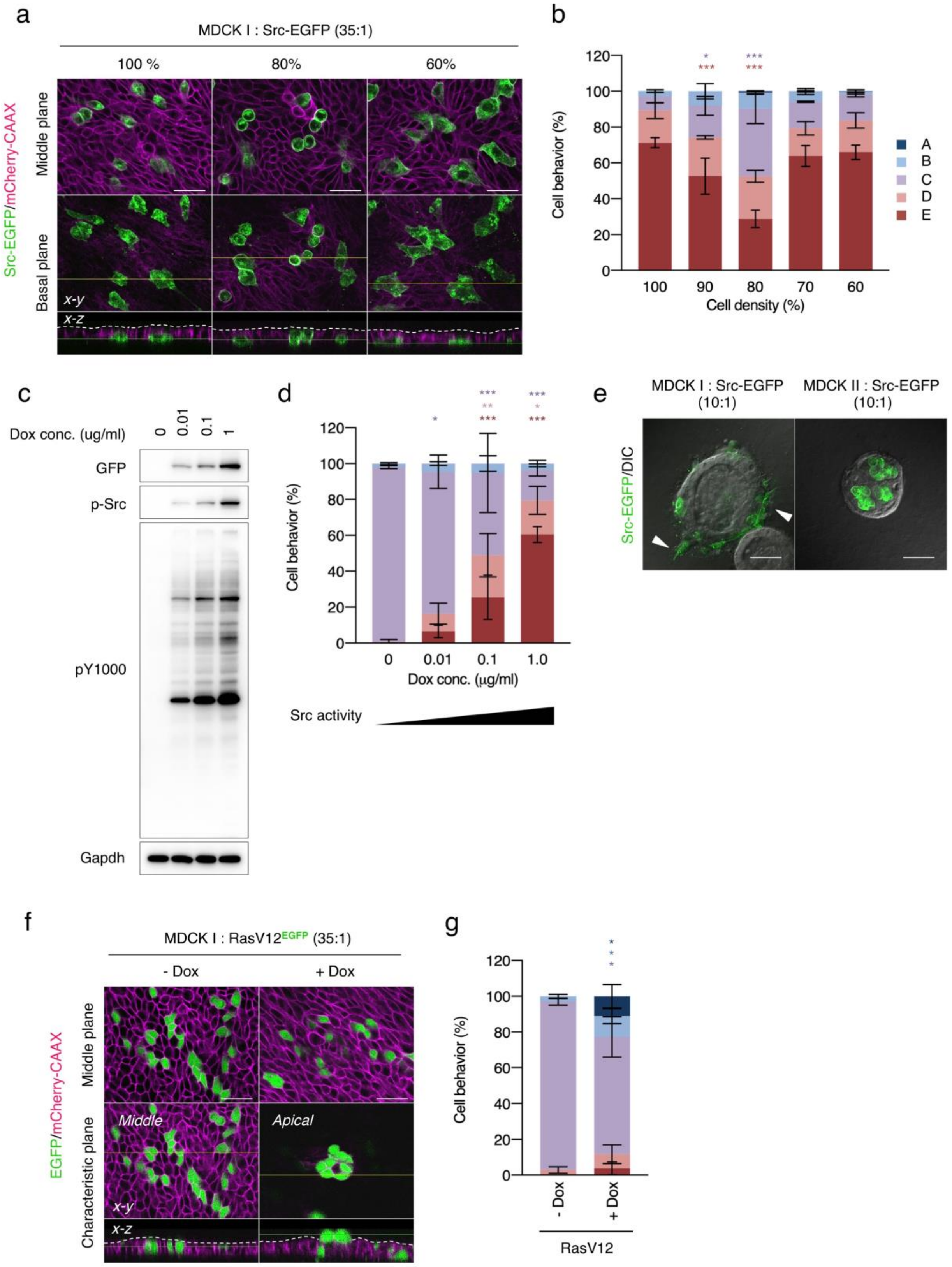
a
MDCK I : Src-EGFP (35:1)
100 %
80%
60%
Src-EGFP/mCherry-CAAX
Middle plane
Basal plane
x-y
x-z
b
Cell behavior (%)
Cell density (%)
A
B
C
D
E
c
Dox conc. (ug/ml)
0
0.01
0.1
1
GFP
p-Src
pY1000
Gapdh
d
Cell behavior (%)
Dox conc. (μg/ml)
Src activity
e
MDCK I : Src-EGFP (10:1)
MDCK II : Src-EGFP (10:1)
Src-EGFP/DIC
f
MDCK I : RasV12EGFP (35:1)
- Dox
+ Dox
EGFP/mCherry-CAAX
Middle plane
Characteristic plane
Middle
Apical
x-y
x-z
g
Cell behavior (%)
- Dox
+ Dox
RasV12

**Supplementary Figure 1. Src-transformed cells delaminate into basal side of epithelial cell layer**

(**a**, **b**) MDCK type I cells harbouring Src-EGFP were mixed with wild-type cells at a ratio of 35:1 and grown on collagen matrix in the presence or absence of 1 μg/ml Dox for 24 h. Cell density of confluent cultures set 100%, and cell behaviour was analysed at each cell density. (**c**, **d**) Type I cells harbouring Src-EGFP were grown in the presence of indicated concentration of Dox for 24 h. Cell lysates were subjected to immunoblotting analysis using the indicated antibodies. (**e**) Type I or type II cells harbouring Src-EGFP were mixed with wild-type cells at a ratio of 10:1, the mosaic cysts were grown in collagen matrix in the presence or absence of 1 μg/ml Dox for 2 days. (**f**, **g**) Type I cells harbouring RasV12 and EGFP were mixed with wild-type cells at a ratio of 35:1 and grown on collagen matrix in the presence or absence of 1 μg/ml Dox for 24 h. Cell behaviour was assessed by using the criteria (Fig. 1c). The mean ratios ± SDs were obtained from three independent experiments. *, $p < 0.05$; **, $p < 0.01$; ***, $p < 0.001$; two-way ANOVA (b, d) or two-tailed *t*-test (e, g) was calculated. The scale bars indicate 50 μm.

Supplementary Figure 2

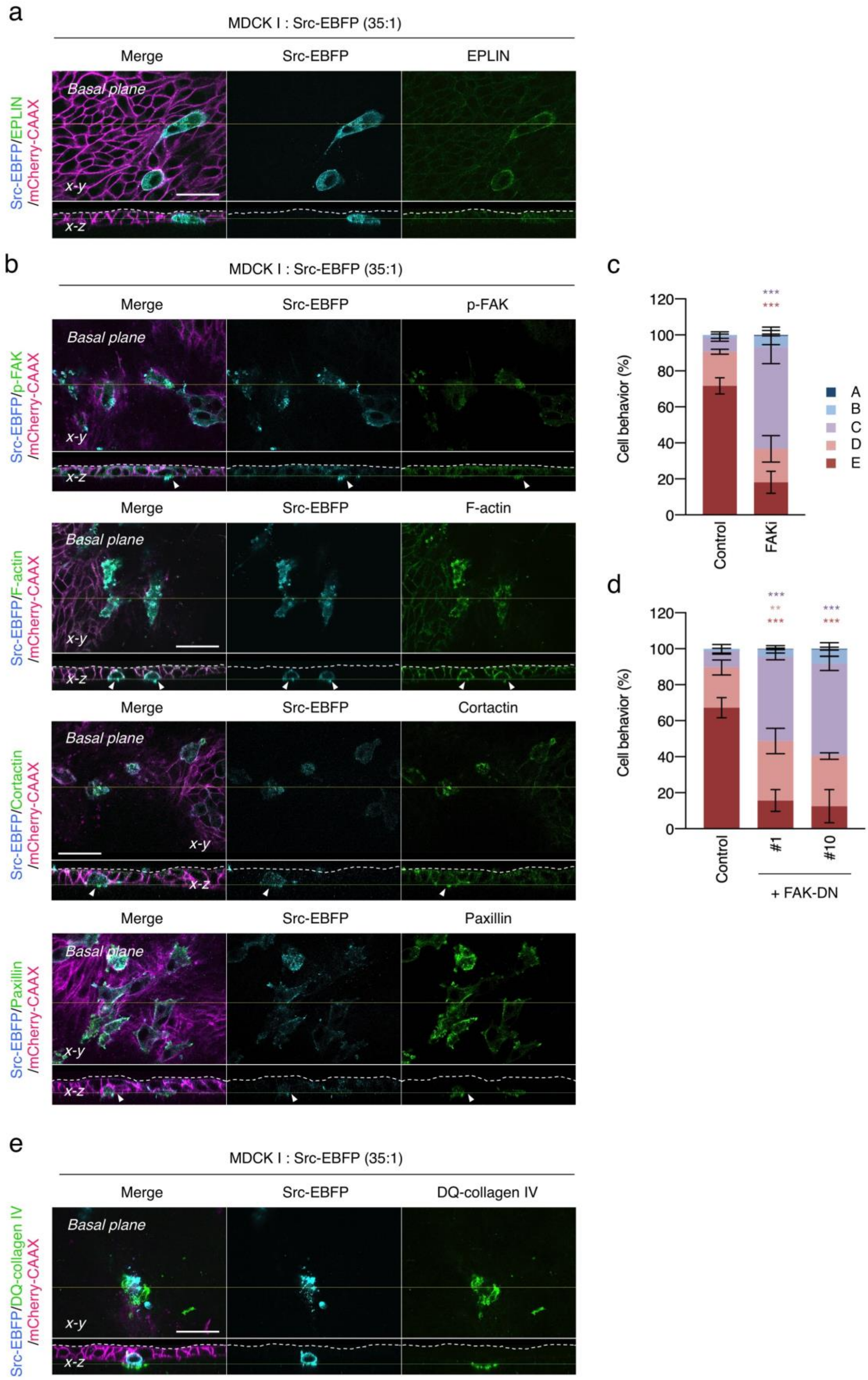

**Supplementary Figure 2. Src-transformed cells delaminate with maturation of focal adhesions**

(**a**, **b**) MDCK type I cells harbouring Src-EBFP were mixed with wild-type cells at a ratio of 35:1 and grown in the presence of Dox for 24 h. After fixation, EPLIN (a), FAK pY576, cortactin, and paxillin (b) were visualized with specific antibodies and Alexa Fluor 488-conjugated secondary antibody. Filamentous actin was stained with Alexa488-conjugated phalloidin. (**c**) Type I cells harbouring Src-EGFP were mixed with wild-type cells at a ratio of 35:1 and preincubated with 200 nM FAK inhibitor 14 (FAKi) for 2 h and then incubated with Dox for 24 h. (**d**) Type I clones harbouring Src-EGFP and FAK-DN were mixed with wild-type cells at a ratio of 35:1 and incubated with Dox for 24 h. Cell behaviour was assessed by using the criteria (Fig. 1c). The mean ratios ± SDs were obtained from three independent experiments. **, $p < 0.01$; ***, $p < 0.001$; two-way ANOVA was calculated compared to non-treated control (c) or mock-transfected cells (d). (**e**) Type I cells harbouring Src-EBFP were mixed with wild-type cells at a ratio of 35:1 and grown on DQ-collagen-containing matrix in the presence of Dox for 24 h. DQ fluorescence represents collagen degradation. The scale bars indicate 50 μm.

Supplementary Figure 3

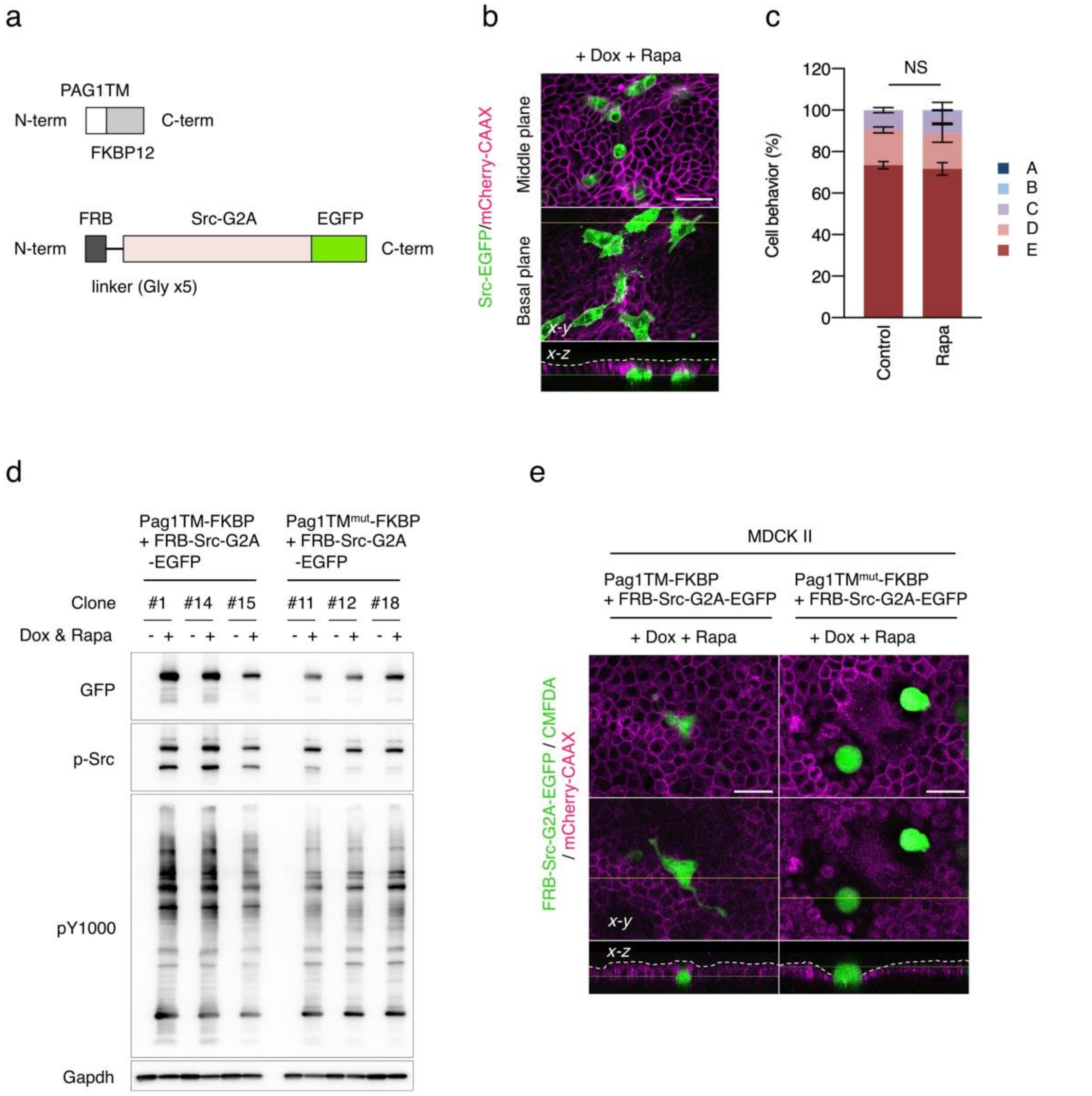

**Supplementary Figure 3. Lipid raft localisation of Src is critical determinant for basal delamination**

(**a**) Schematic diagram of Pag1TM-FKBP (or Pag1TM$^{mut}$-FKBP) and FRB-Src-G2A-EGFP constructs. FKBP fused to the C-terminus of the Pag1/Cbp transmembrane domain with or without the palmitoylation motif (Pag1TM-FKBP or Pag1TM$^{mut}$-KFBP) was stably expressed in MDCK type I cells. In contrast, the FRB domain fused to the N-terminus of the Src-EGFP (FRB-Src-G2A-EGFP) cytosolic form was expressed by a Dox-inducible system. In this system, FRB-Src-G2A-EGFP expressed in the cytosol by Dox treatment was translocated to either lipid raft or non-raft compartments by treatment with rapamycin (See also, Fig. 3a). (**b**, **c**) Type I cells harbouring Src-EGFP were mixed with wild-type cells at a ratio of 35:1 and preincubated with 100 nM rapamycin (Rapa) for 2 h and then incubated with 1 µg/ml Dox for 24 h. Cell behaviour was assessed by using the criteria (Fig. 1c). The mean ratios ± SDs were obtained from three independent experiments. NS, not significantly different; two-way ANOVA was calculated compared to non-treated control. (**d**) Type I cells harbouring FRB-Src-G2A-EGFP and Pag1TM-FKBP or Pag1TM$^{mut}$-FKBP were incubated in the presence of 1 µg/ml Dox and 100 nM Rapa for 24 h. Lysates from these cells were analysed by immunoblotting using indicated antibodies. (**e**) Type II cells harbouring FRB-Src-G2A-EGFP and Pag1TM-FKBP or Pag1TM$^{mut}$-FKBP were mixed with wild-type cells at a ratio of 35:1 and incubated in the presence of Dox and 100 nM Rapa for 24 h. The scale bars indicate 50 µm.

Supplementary Figure 4

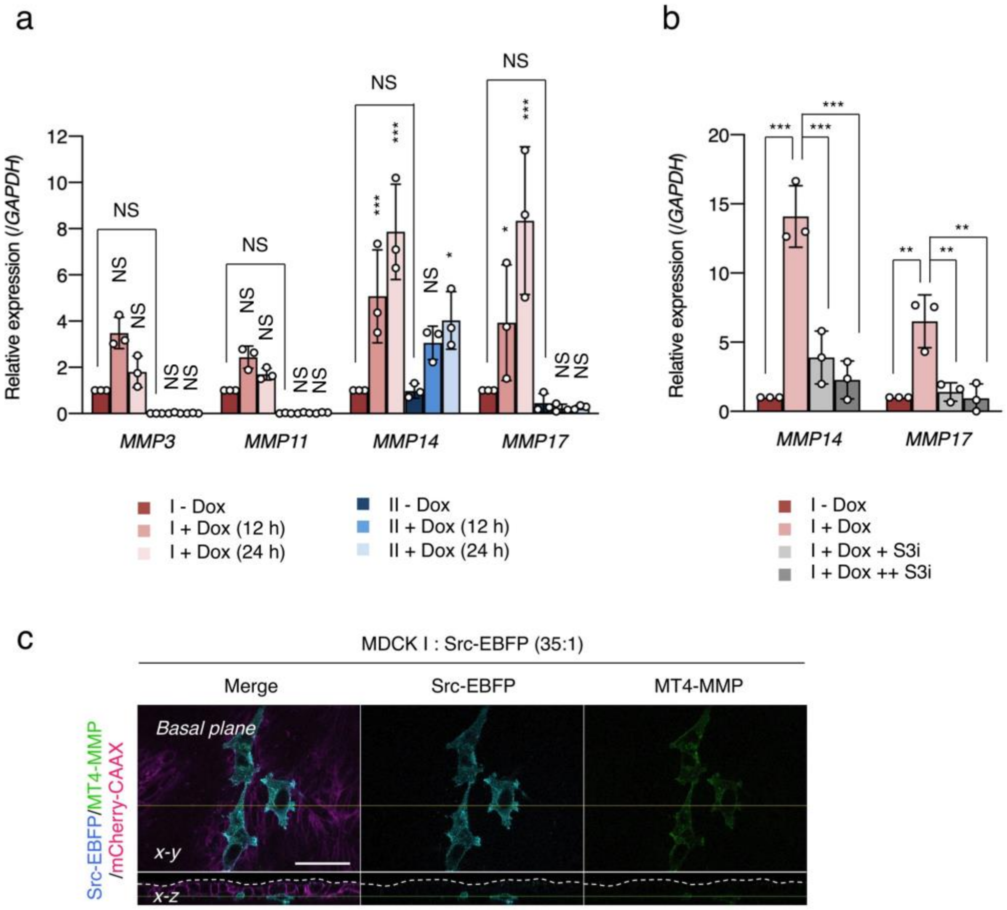

**Supplementary Figure 4. Expression of MMP genes is increased by the Src-STAT3 signalling**

(**a**) MDCK type I and II harbouring Src-EGFP were incubated in the presence of Dox for indicated time periods. Expression of MMP mRNAs were analysed by quantitative PCR analysis. (**b**) Type I cells harbouring Src-EGFP were mixed with wild-type cells at a ratio of 35:1 and preincubated with S3i-201 (+ S3i, 50 μM; ++ S3i, 100 μM) for 2 h and then incubated with Dox for 24 h. Expression of MMP mRNAs were analysed by quantitative PCR analysis. The mean ratios ± SDs were obtained from three independent experiments. *, $p < 0.05$; **, $p < 0.01$; ***, $p < 0.001$; NS, not significantly different; two-way ANOVA was calculated compared to non-treated cells. (**c**) Type I cells harbouring Src-EBFP were mixed with wild-type cells at a ratio of 35:1 and grown in the presence of Dox for 24 h. After fixation, MT4-MMP was visualized with specific antibodies and Alexa Fluor 488-conjugated secondary antibody.

### Supplementary Figure 5

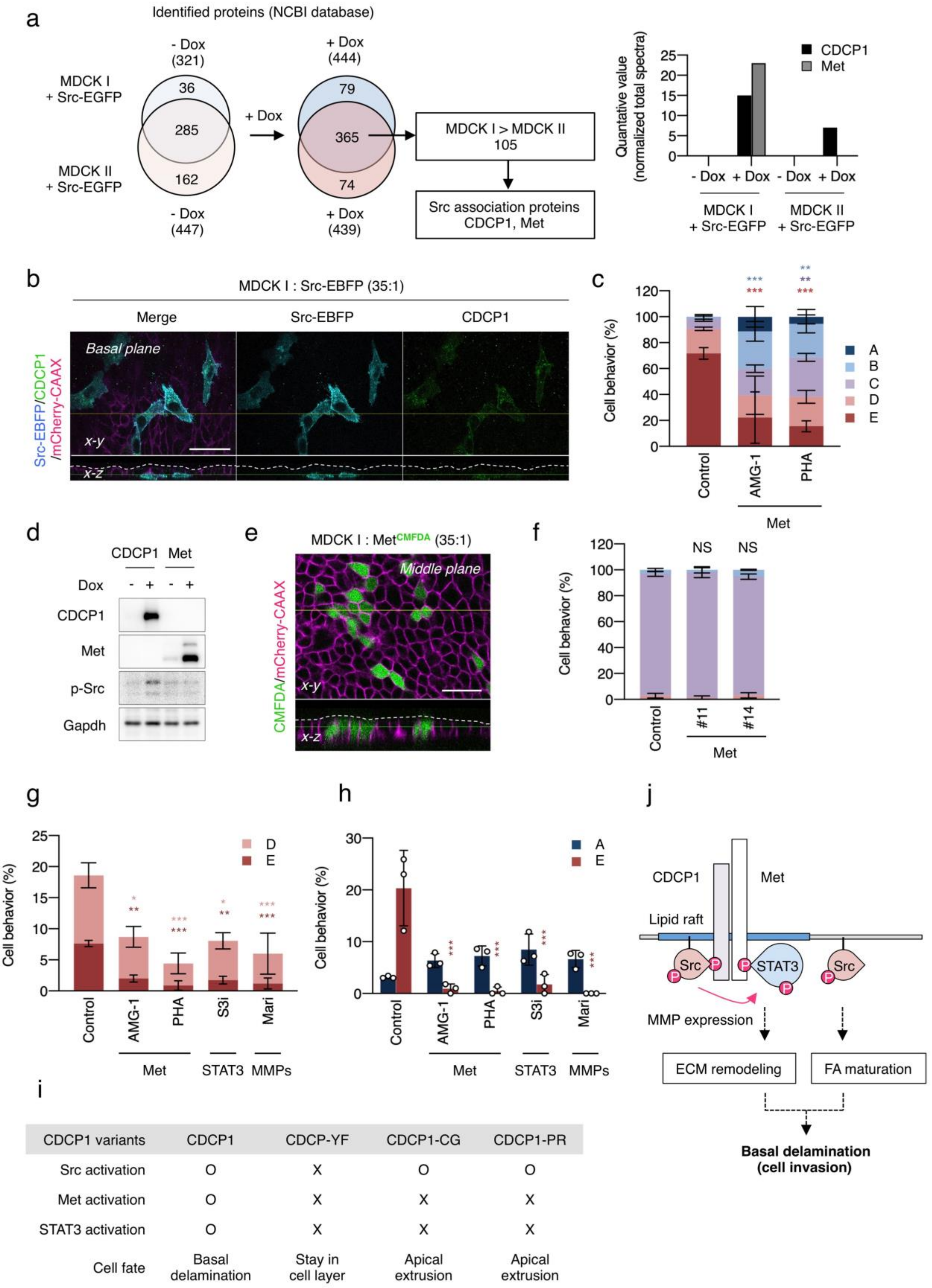

| CDCP1 variants | CDCP1 | CDCP-YF | CDCP1-CG | CDCP1-PR |
|---|---|---|---|---|
| Src activation | O | X | O | O |
| Met activation | O | X | X | X |
| STAT3 activation | O | X | X | X |
| Cell fate | Basal delamination | Stay in cell layer | Apical extrusion | Apical extrusion |

**Supplementary Figure 5. CDCP1 is a lipid raft-localised Src recruiting protein**

(**a**) Schematic flow diagram of identification of Src-recruiting protein candidates. MDCK type I and II harbouring Src-EGFP were grown in the presence or absence of Dox for 24 h. Lysates from these cells were subjected to the DRM separation experiment. Aliquots of the DRM fractions were analysed by silver staining. All proteins within polyacrylamide gel were trypsinised and subjected mass spectrometry. Proteins were identified using the NCBI database and the number of identified proteins is depicted. Right graph indicated quantified value of identified proteins. See also Supplementary Table 3. (**b**) Type I cells harbouring Src-EBFP were mixed with wild-type cells at a ratio of 35:1 and grown in the presence of Dox for 24 h. After fixation, CDCP1 was visualized with specific antibodies and Alexa Fluor 488-conjugated secondary antibody. (**c**) Type I cells harbouring Src-EGFP were mixed with wild-type cells at a ratio of 35:1 and preincubated with 200 nM AMG-1 of 400 nM PHA-665752 (PHA) for 2 h and then incubated with Dox for 24 h. (**d**) Type I cells harbouring CDCP1-EGFP or Met were grown in the presence or absence of Dox for 24 h. Cell lysates were subjected to immunoblotting analysis using the indicated antibodies. (**e, f**) Type I clones harbouring Met were stained with CMFDA and mixed with wild-type cells at a ratio of 35:1 and then incubated with Dox for 24 h. (**g**) Type I cells harbouring CDCP1-EGFP were mixed with wild-type cells at a ratio of 35:1 and preincubated with 200 nM AMG-1, 400 nM PHA-665752 (PHA), 100 μM S3i-201 (S3i), 20 μM stattic (Sta), or 20 μM marimastat (Mari) for 2 h and then incubated with Dox for 24 h. (**h**) Type I cells harbouring CDCP1-EGFP were mixed with wild-type cells at a ratio of 8:1, and the mosaic spheroid were preincubated with the 200 nM AMG-1, 400 nM PHA-665752 (PHA), 100 μM S3i-201 (S3i), 20 μM stattic (Sta), or 20 μM marimastat (Mari) for 2 h and then incubated with Dox for 2 days. Cell behaviour was assessed by using the criteria (Fig.1c). The mean ratios ± SDs were obtained from three independent experiments. *, $p < 0.05$; **, $p < 0.01$; ***, $p < 0.001$; NS, not significantly different; two-way ANOVA was calculated compared to non-treated cells (c, i) or mock-transfected cells (e, h). The scale bars indicate 50 μm. (**i**) A summary table of CDCP1 variants-mediated phenomena and cell delamination. Src activation induces bidirectional cell delamination. Basal delamination is occurred when activation of Met-STAT3 signalling. (**j**) Schematic model of Src-induced cell delamination. Activated Src is recruited and trapped in lipid rafts by CDCP1. Src phosphorylates STAT3 through CDCP1-Met association in lipid rafts. Activated STAT3 enhances cell migration and expression of MMPs in which induces remodelling of ECM. Simultaneously, activated Src induces maturation of focal adhesions (FA) in which induce adhesion to ECM. These Src-mediated events promote basal delamination in an integrated manner against apical extrusion.

Supplementary Figure 6

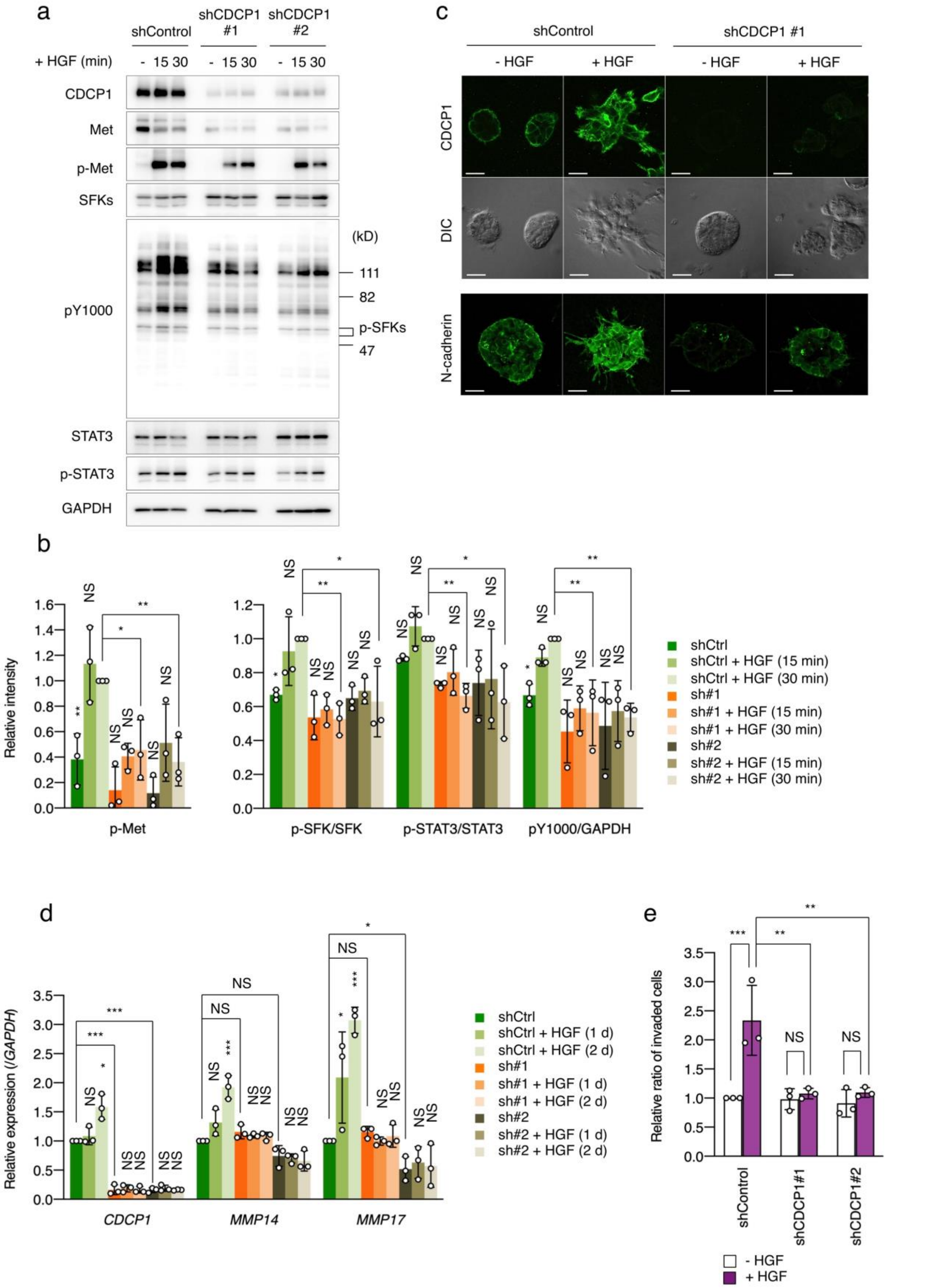

**Supplementary Figure 6. CDCP1 is required for HGF-mediated EMT progression**

(**a, b**) shControl or shCDCP1 expressing A498 cells were incubated in the presence of 50 ng/ml HGF for indicated time periods. Phosphorylation levels were analysed by western blot analysis. The mean ratios ± SDs were obtained from three independent experiments. (**c**) shControl or shCDCP1 expressing A498 spheroids were incubated in the presence of 50 ng/ml HGF for 2 days. After fixation, CDCP1 and N-cadherin were visualized with specific antibodies and Alexa Fluor 488-conjugated secondary antibody. The scale bars indicate 50 µm. (**d**) shControl or shCDCP1 expressing A498 cells were incubated in the presence of 50 ng/ml HGF for 1 or 2 days. Expression of indicated mRNAs were analysed by quantitative PCR analysis. The mean ratios ± SDs were obtained from three independent experiments. (**e**) shControl or shCDCP1 expressing A498 cells were subjected to matrigel invasion assay in the presence of 50 ng/ml HGF. The mean ratios ± SDs were obtained from three independent experiments. *, $p < 0.05$; **, $p < 0.01$; ***, $p < 0.001$; NS, not significantly different; two-way ANOVA was calculated compared to HGF-treated (b, e) or -untreated shControl cells (d).

Supplementary Figure 7

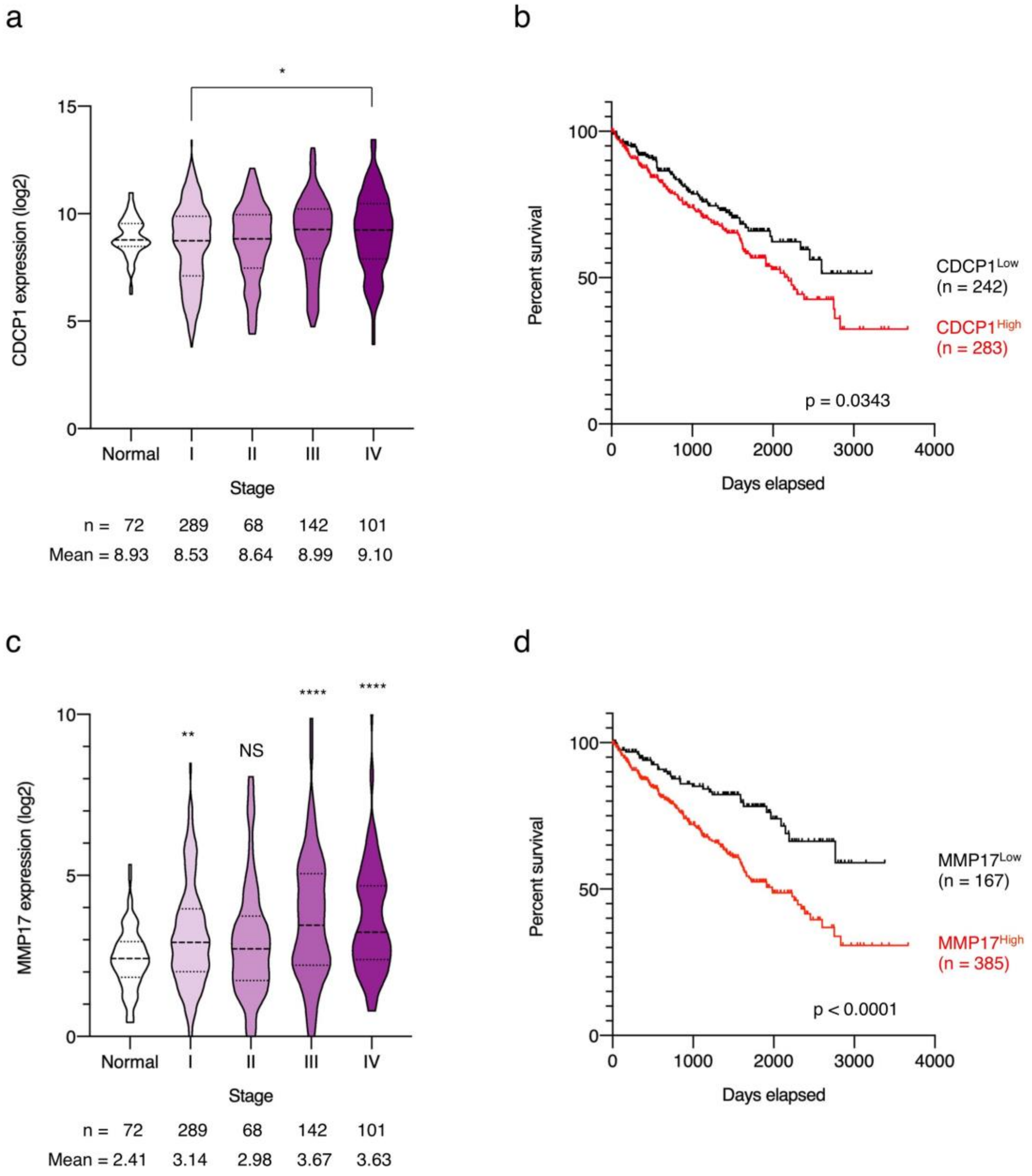

**Supplementary Figure 7. Expression level of CDCP1 and MMP17 is associated with poor prognosis of renal cancer patients**

(**a, c**) mRNA expression of CDCP1 (a) and MMP17 (c) was analysed by using cohort of kidney clear cell carcinoma patients (n = 600) from The Cancer Genome Atlas. Expression levels in normal or Stage I-IV patients were separately analysed. **, $p < 0.01$; ***, $P < 0.001$; ****, $p < 0.0001$; NS, not significantly different; one-way ANOVA was calculated compared to Stage I tumour (a) or normal tissue (b). The mean expression level was calculated. (**b, d**) Kaplan-Meier survival curves of kidney clear cell carcinoma patients were constructed using mean expression (high vs low) and analysed with the log-rank test.

**Supplementary Table 1. Inhibitor screening data.**

| Category | Compound | Inhibition |
|---|---|---|
| Met | PHA665752 | ++ |
| Met | Met/Ron inhibitor | + |
| ALK | A83-01 | - |
| Notch | DAPT | - |
| Src | Dasatinib | +++ |
| FAK | FAK inhibitor 14 | ++ |
| JAK | JAK Inhibitor I | - |
| JAK | Ruxolitinib | - |
| STAT3 | WP1066 | + |
| STAT3 | 5,15-DPP | + |
| STAT3 | Stattic | + |
| STAT3 | S3i-201 | ++ |
| PI3K | LY294002 | ++ |
| p70 S6K | Rapamycin | - |
| GSK3 | GSK-3 inhibitor IX | ++ |
| GSK3 | 1-Azakenpaullone | ++ |
| GSK3 | TWS119 | ++ |
| GSK3 | CT99021 | ++ |
| MEK | U0126 | ++ |
| ROCK | H-1152 | +++ |
| ROCK | Y-27632 | + |
| JNK | SP600125 | - |
| JNK | JNK inhibitor VIII | - |
| YAP-TEAD | Verteporfin | - |
| HIF | Chetomin | - |
| Actin filament | Cytochalasin D | - |
| MMP | GM 6001 | ++ |
| MMP | Marimastat | + |
| Aurora | MLN8237 | + |
| HDAC | Trichostatin A | - |

| | | |
|---|---|---|
| HDAC | Vorinostat | - |
| Serine palmitoyl transferase | Myriocin | + |
| Sphingosine N-acyltransferase | Fumonisin B1 | + |
| Glucosyl ceramide synthase | PDMP | + |
| Fatty acid synthase | C75 | + |
| Fatty acid synthase | Cerulenin | + |
| HMG-CoA reductase | Lovastatin | +++ |
| HMG-CoA reductase | Simvastatin | + |
| Cholesterol | MbCD | +++ |

| | |
|---|---|
| +++ | Strong inhibition (<10%) |
| ++ | Inhibition (<20%) |
| + | Weak inhibition |
| - | No inhibition |

**Supplementary Table 2. Lipid raft-localised proteins in DRM fractions.**

Lipid raft-localised proteins, flotillin and caveolin, in DRM fractions were identified by mass spectrometry using NCBI database and listed.

| Identified Proteins | Normalized total spectra | | | |
|---|---|---|---|---|
| | MDCK I + Src-EGFP | | MDCK II + Src-EGFP | |
| | - Dox | + Dox | - Dox | + Dox |
| flotillin-1 | 29.14 | 34.2 | 37.34 | 31.24 |
| flotillin-2 | 16.65 | 25.4 | 35.47 | 32.08 |
| caveolin-1 | 37.47 | 34.2 | 52.27 | 68.38 |
| caveolin-2 | 12.49 | 9.77 | 6.53 | 15.2 |

**Supplementary Table 3. Proteins enriched after Src overexpression in DRM fractions.**

Proteins in DRM fractions were identified by mass spectrometry using NCBI database. Top 30 of enriched proteins after Src overexpression were listed.

| Identified Proteins | Normalized total spectra | | | |
|---|---|---|---|---|
| | MDCK I + Src-EGFP | | MDCK II + Src-EGFP | |
| | - Dox | + Dox | - Dox | + Dox |
| proto-oncogene tyrosine-protein kinase Src | 11.1 | 232.54 | 10.27 | 46.43 |
| tyrosine-protein kinase Yes | 18.04 | 57.65 | 17.73 | 27.01 |
| focal adhesion kinase 1 | 0 | 35.17 | 0 | 0 |
| urokinase-type plasminogen activator precursor | 0 | 30.29 | 3.73 | 20.26 |
| V-type proton ATPase catalytic subunit A | 18.04 | 47.88 | 100.81 | 109.75 |
| ephrin type-A receptor 2 | 0 | 29.31 | 0.93 | 2.53 |
| arf-GAP with Rho-GAP domain, ANK repeat and PH domain-containing protein 3 | 0 | 28.33 | 0 | 0.84 |
| V-type proton ATPase 116 kDa subunit a isoform 1 | 20.82 | 48.85 | 44.8 | 45.59 |
| tyrosine-protein kinase Fyn | 0 | 27.36 | 0 | 0 |
| semaphorin-7A | 31.92 | 58.62 | 34.54 | 54.03 |
| V-type proton ATPase subunit B | 18.04 | 43.97 | 81.2 | 61.63 |
| solute carrier family 2, facilitated glucose transporter member 1 | 22.2 | 44.94 | 37.34 | 51.5 |
| trifunctional enzyme subunit alpha, mitochondrial | 0 | 22.47 | 38.27 | 17.73 |
| hepatocyte growth factor receptor | 0 | 22.47 | 0 | 0 |
| desmoglein-2 | 26.37 | 46.9 | 55.07 | 71.76 |
| 78 kDa glucose-regulated protein | 0 | 19.54 | 11.2 | 19.42 |
| V-type proton ATPase subunit B | 0 | 18.56 | 54.14 | 41.37 |
| desmoglein-3 precursor | 48.57 | 65.46 | 27.07 | 57.41 |
| CD109 antigen | 34.69 | 50.81 | 59.74 | 70.91 |
| urokinase plasminogen activator surface receptor | 4.16 | 19.54 | 0 | 14.35 |
| lamin | 8.33 | 23.45 | 21.47 | 21.95 |
| CUB domain-containing protein 1 | 0 | 14.66 | 0 | 6.75 |
| galectin-3-binding protein | 0 | 14.66 | 0 | 1.69 |
| plakophilin-2 | 0 | 13.68 | 14 | 25.33 |
| protein EFR3 homolog A | 16.65 | 30.29 | 15.87 | 12.66 |

| | | | | |
|---|---|---|---|---|
| V-type proton ATPase 116 kDa subunit a | 2.78 | 15.63 | 19.6 | 17.73 |
| tax1-binding protein 1 | 0 | 12.7 | 3.73 | 10.97 |
| laminin subunit beta-3 | 0 | 12.7 | 0 | 9.29 |
| sodium/potassium-transporting ATPase subunit alpha-1 precursor | 8.33 | 20.52 | 14 | 14.35 |
| catenin delta-1 | 24.98 | 37.13 | 5.6 | 11.82 |

**Supplementary Table 4. Primer list of quantitative real-time PCR**

| Genes | Forward primer (5'-3') | Reverse primer (5'-3') |
|---|---|---|
| Canis MMP3 | tgcagttagagatcacggagac | gggtaggcatgacccaaaa |
| Canis MMP11 | tgggataaaacggacctcacc | cacgaccctcatgcacct |
| Canis MMP14 | ctcgccccagtcattctc | acgcgcagaccatagaactt |
| Canis MMP17 | ctctatcatgcagccctactacc | acacagactcccgcacac |
| Canis GAPDH | gattgtcagcaatgcctcct | ggtcatggatgactttggcta |
| Homo sapiens CDCP1 | aggaaggaggagcgggtt | tggactttgggcatcttggt |
| Homo sapiens MMP14 | gttctggcgggtgaggaataa | ctcgtaggcagtgttgatgga |
| Homo sapiens MMP17 | ctgtacggtgtgcgggag | tgagtgctgcatctgtggg |
| Homo sapiens GAPDH | agtccactggcgtcttcac | tcttgaggctgttgtcatacttct |
